# Multi-protein Bridging Factor 1(Mbf1), Rps3 and Asc1 prevent stalled ribosomes from frameshifting

**DOI:** 10.1101/366344

**Authors:** Jiyu Wang, Jie Zhou, Qidi Yang, Elizabeth J. Grayhack

**Author notes:** **Abbreviations:** helix-turn-helix (HTH) Ribosome Quality Control (RQC).

## Abstract

Stalled ribosomes in bacteria frameshift, but stalled ribosomes in eukaryotes do not frameshift and abort translation, suggesting that eukaryote-specific mechanisms might prevent frameshifting. We show that the conserved eukaryotic/archaeal protein Mbf1 acts with ribosomal proteins Rps3/uS3 and eukaryotic Asc1/RACK1 to prevent frameshifting at inhibitory CGA-CGA codon pairs in *Saccharomyces cerevisiae*. Mutations in *RPS3* that allow frameshifting implicate eukaryotic conserved residues near the mRNA entry site. Mbf1 and Rps3 cooperate to maintain the reading frame of stalled ribosomes, while Asc1 mediates distinct events that result in aborted translation. Frameshifting occurs through a +1 shift with a CGA codon in the P site and involves competition between codons entering the A site, implying that the wobble interaction of the P site codon destabilizes translation elongation. Thus, eukaryotes have evolved unique mechanisms involving both a universally conserved ribosome component and two eukaryotic-specific proteins to maintain the reading frame at ribosome stalls.

## INTRODUCTION

Accurate translation of mRNA into protein depends upon precise, repetitive three base translocation of the ribosome to maintain the correct reading frame throughout a coding sequence. Reading frame maintenance is challenging because multiple movements of the tRNAs and mRNA as well as conformational changes within the ribosome itself are required to complete a single elongation cycle (Noller et al., 2017). For instance, the tRNA acceptor stems move within the large subunit during formation of the hybrid state, while the joining of EF-G-GTP (eEF2 in eukaryotes) results in additional movement of tRNA (Brilot et al., 2013, Ramrath et al., 2013, Zhou et al., 2014), and finally completion of translocation, driven by Pi release, requires additional movements (Noller et al., 2017). (Brilot et al., 2013, Ramrath et al., 2013, Zhou et al., 2014). To accomplish this cycle, many interactions between the tRNAs and ribosome are disrupted, and new interactions are created, but the relative position of the tRNA anticodon to the mRNA codon must be maintained throughout all of these events (Noller et al., 2017, Dever et al., 2018, Rodnina, 2018). Thus, it is critical that mechanisms exist to prevent slippage during these transitions.

Reading frame maintenance is facilitated by structures within the ribosome as well as by tRNA modifications. Structural features that contribute to reading frame maintenance, inferred from analysis of prokaryotic translation intermediates, include a swivel of the 30S head relative to the 30S body to form a contracted mRNA tunnel downstream of the A site prior to translocation (Jenner et al., 2010, Schuwirth et al., 2005). In addition, during translocation, two conserved bases in the 16S rRNA intercalate into different positions of mRNA to prevent slippage (Zhou et al., 2013) while domain IV of EF-G contacts both the codon-anticodon in the A/P site and the 16S rRNA, likely coupling mRNA and tRNA movement (Ramrath et al., 2013, Zhou et al., 2014). tRNA modifications within the anticodon loop also assist in reading frame maintenance, inferred both from genetic and structural analyses. Mutants that affect several such modifications in both bacteria and eukaryotes result in increased frameshifting (Atkins and Bjork, 2009, Jager et al., 2013, Tukenmez et al., 2015, Urbonavicius et al., 2001, Waas et al., 2007). Moreover, a cross-strand base stacking interaction between a modified ms^2^i^6^A37 in an *E. coli* tRNA^Phe^ and the mRNA codon is proposed to prevent slippage of P site tRNA on the mRNA (Jenner et al., 2010). Thus, a number of mechanisms exist to prevent loss of reading frame.

Nevertheless, ribosomes do move into alternative reading frames in response to specific sequences and structures in mRNA (Atkins and Bjork, 2009, Dever et al., 2018, Dinman, 2012). The existence of such events has implied that ribosomal plasticity with respect to reading frame movement is an integral function of the translation machinery. The common feature of all frameshifting events in bacteria to humans is that the ribosome stalls (Dever et al., 2018). The stall can be mediated in several different ways, by combined effects of the A and P site codons (Farabaugh et al., 2006, Gamble et al., 2016) or by the presence of the downstream structures or an upstream Shine-Dalgarno sequence in bacteria (Caliskan et al., 2014, Dinman, 2012). Analysis of programmed frameshifting indicates that there are frequently requirements for additional sequences or protein factors to mediate efficient frameshifting (Atkins and Bjork, 2009, Dinman, 2012). For instance, +1 programmed frameshifting events are frequently enhanced by stimulatory sequences, although the role of these sequences is not always clear (Guarraia et al., 2007, Taliaferro and Farabaugh, 2007).

The identification of mutants that either affect programmed frameshifting or suppress frameshift mutations has pointed to four key factors in reading frame maintenance. First, mutations of ribosomal proteins, particularly those that contact the P site tRNA can cause increased frameshifting. In bacteria, frameshifting mutations are suppressed by deletions within the C terminal domain of ribosomal protein uS9, which contacts the P site tRNA anticodon loop (Jager et al., 2013). In the yeast *Saccharomyces cerevisiae*, programmed frameshifting of the L-A virus is affected by mutations in 5S rRNA or its interactors uL18 or uL5, that also contact the P site tRNA (Meskauskas and Dinman, 2001, Rhodin and Dinman, 2010, Smith et al., 2001). Frameshifting mutations are also suppressed by a mutation in the yeast *RPS3*, although this mutation does not affect a tRNA contact (Hendrick et al., 2001). Second, mutations in the basal translation machinery can also affect frameshifting. For instance, frameshifting mutations are suppressed by mutations in both EF-1α, which delivers tRNA to the ribosome (Sandbaken and Culbertson, 1988), and in *SUP35*, encoding the translation termination factor eRF3 (Wilson and Culbertson, 1988). Third, miRNAs can affect the efficiency of programmed frameshifting, for instance at *CCR5* in humans (Belew et al., 2014). Fourth, mutations that affect proteins with previously unknown functions in translation can alter either programmed frameshifting or suppress frameshifting mutations. For instance, in yeast, frameshifting mutations are suppressed by mutations in *MBF1*, encoding Multiprotein Bridging Factor 1 (Hendrick et al., 2001), or in *EBS1* (Ford et al., 2006), while in the porcine virus PRRSV, the RNA binding protein nsp1β stimulates both −1 and −2 frameshifting events (Li et al., 2014). Thus, reading frame maintenance is modulated by ribosomal components, many of which contact the tRNAs, as well as by extra-ribosomal proteins and miRNAs. However, the roles of many of these proteins are not understood.

We set out to work out the mechanisms that maintain reading frame when eukaryotic ribosomes encounter a stall, the common feature of all frameshifting events. In bacteria, ribosome stalls due to limited availability or functionality of tRNA seem to suffice to cause frameshifting (Atkins 2005, Hayes 2011). However, in eukaryotes, the rescue of stalled ribosomes by frameshifting is not observed. In wild type yeast, ribosomes stall at CGA codon repeats, which inhibit translation due to wobble decoding of CGA by its native tRNA^Arg(ICG)^ (Letzring et al., 2010, Letzring et al., 2013), but do not frameshift (Wolf and Grayhack, 2015). Instead, eukaryotes have evolved new pathways to regulate inefficient translation events, such as the Ribosome Quality Control (RQC) pathway, in which these stalled ribosomes undergo ubiquitination of ribosomal proteins, followed by dissociation of the subunits, and recruitment of the RQC Complex, which mediates CAT tailing and degradation of the nascent polypeptide (Brandman and Hegde, 2016, Brandman et al., 2012, Joazeiro, 2017, Juszkiewicz and Hegde, 2017, Matsuo et al., 2017, Simms et al., 2017, Sundaramoorthy et al., 2017, Shen et al., 2015). The ribosomal protein Asc1/RACK1 mediates these events (Brandman et al., 2012, Kuroha et al., 2010); in the absence of Asc1, ribosomes continue translation through CGA codon repeats more efficiently, but also undergo substantial frameshifting at these repeats (Wolf and Grayhack, 2015). However, Asc1 sits on the outside of the ribosome at the mRNA exit tunnel and likely functions as scaffold for recruitment of other proteins, such as the E3 ubiquitin ligase Hel2/mammalian ZNF598 and Slh1 (Kostova et al., 2017, Matsuo et al., 2017, Sitron et al., 2017). Based on the location of Asc1 and the precedent that Asc1 recruits other proteins to abort translation, we considered it likely that Asc1 cooperates with additional proteins to mediate reading frame maintenance at CGA codon repeats and set out to find such factors.

Here, we provide evidence that the Multiprotein Bridging Factor 1 (Mbf1) and ribosomal protein Rps3 work together to prevent translational slippage at CGA codon repeats. Frameshifting is caused by inactivation of *MBF1*, and by mutations of amino acids in Rps3 located on an exposed surface of the protein near the mRNA entry site. Frameshifting in *RPS3* mutants is suppressed by additional copies of the *MBF1* gene. We provide evidence that Asc1 mediates a distinct, but related function to that of Mbf1, acting to abort translation of stalled ribosomes, which in turn reduces frameshifting. Mbf1 and Asc1 synergistically prevent frameshifting at the seven most slowly translated codon pairs in yeast, all of which are codon pairs that inhibit translation relative to their synonymous optimal pairs (Gamble et al., 2016). We examine the precise frameshift at one of these inhibitory pairs, CGA-CGG, purifying the frameshifted polypeptide, followed by analysis with mass spectrometry. We find that frameshifting occurs in the +1 direction at the CGA codon and moreover, that frameshifting is modulated by the competition between the in-frame and +1 frame tRNAs.

## RESULTS

### *MBF1* (Multiprotein-Bridging Factor 1) prevents frameshifting at CGA codon repeats

We considered it likely that proteins other than Asc1 worked to prevent frameshifting at CGA codon repeats for two reasons. First, Asc1 binds on the outside of the ribosome, not in the decoding center (Rabl et al., 2011), and thus is not positioned in any obvious way to assist with reading frame maintenance. Second, Asc1 recruits other proteins, Hel2 and Slh1, to bring about aborted translation (Brandman and Hegde, 2016, Joazeiro, 2017), and thus is likely to work with other proteins in reading frame maintenance. We note that Hel2 is not involved (Wolf and Grayhack, 2015). Thus, we set out to identify genes responsible for reading frame maintenance at CGA codon repeats.

To isolate mutants that frameshift due to translation of CGA codon repeats, we set up a selection in which expression of the *URA3* gene depended upon a +1 frameshift due to the presence of 6 adjacent CGA codons. The native *URA3* gene was placed in the +1 reading frame downstream of an N-terminal domain of *GLN4* encoding amino acids 1-99 (*GLN4*_(1-99)_), followed by 6 CGA codons and one additional nucleotide upstream of the *URA3* coding region (Fig. 1A). Thus, this strain exhibits an Ura^-^ phenotype, due to the low levels of frameshifting in an otherwise wild-type background. As a secondary screen for frameshifting mutants due to CGA codon repeats, we integrated a modified version of the RNA-ID reporter with *GLN4*_(1-99)_ followed by 4 CGA codons and one additional nucleotide upstream of the *GFP* coding region into the *ADE2* locus (Dean and Grayhack, 2012, Wolf and Grayhack, 2015). Thus, GFP expression was dependent upon frameshifting efficiency (Fig. 1A). To avoid obtaining mutations in the *ASC1* gene, the selection strain also contained a second copy of the *ASC1* gene on a plasmid. (Fig. 1A). We selected Ura^+^ mutants from 40 independent cultures each of *MAT****a*** and *MATα* parents and then analyzed three Ura^+^ mutants from each culture by flow cytometry to measure GFP and RFP expression. Most mutants (60% of *MATα* mutants and 80% of *MAT****a*** mutants) showed elevated expression of GFP (Fig. 1B), and we studied those that exhibited relatively high levels of frameshifting, >30% of that in an *asc1*Δ mutant. Most mutants (43 of 48 examined) were recessive and mapped to single complementation group, based their growth on media lacking uracil (Fig. 1-figure supplement 1A), although four dominant mutants were also identified.

**Figure 1.**
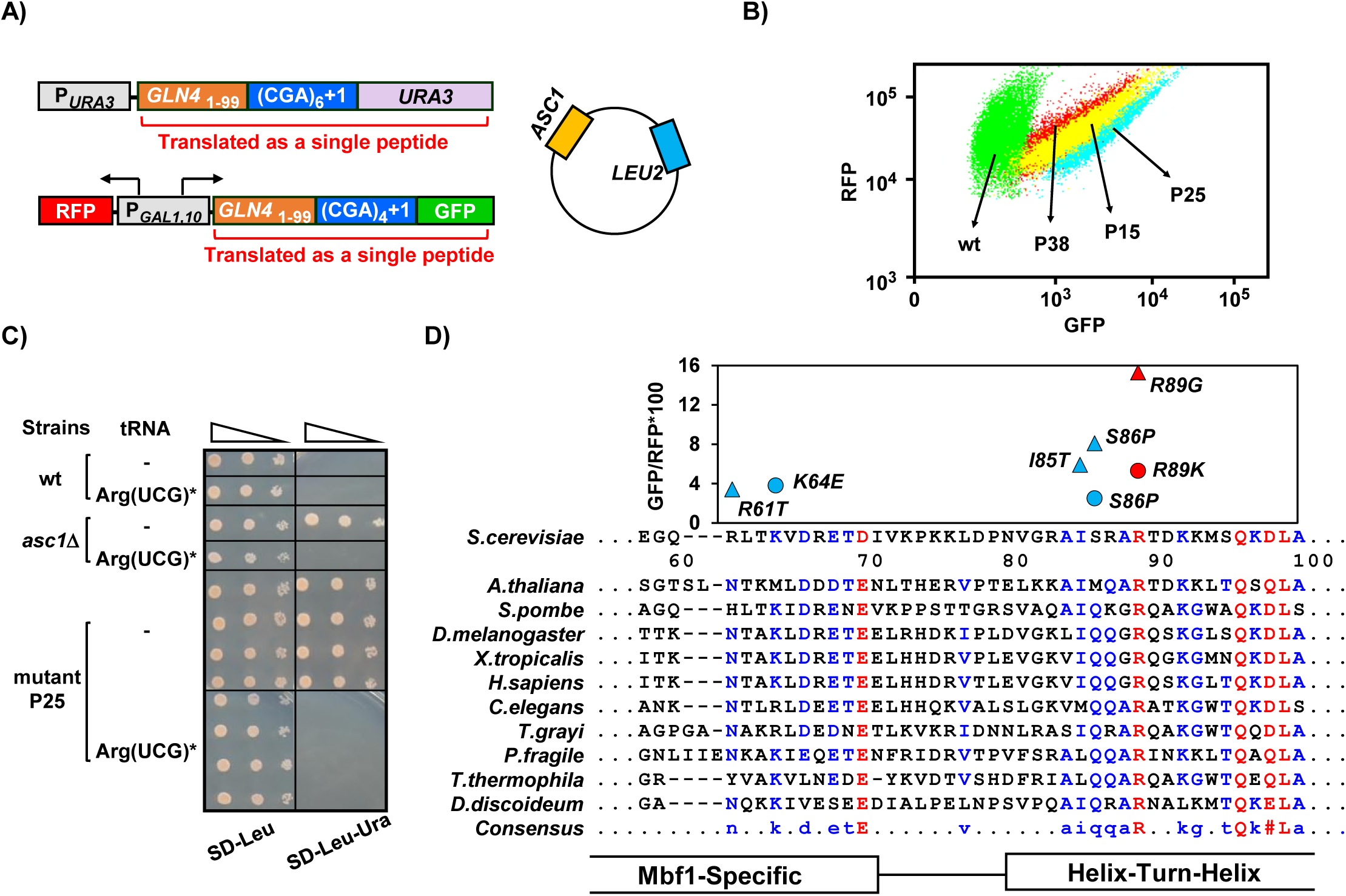
*MBF1* (Multiprotein-Bridging Factor 1) prevents frameshifting at CGA codon repeats. **(A)** Schematic of selection for mutants that frameshift at CGA codon repeats. The indicated CGA codon repeats plus one extra nucleotide were inserted upstream of the *URA3* and *GFP* coding region, resulting in an Ura^-^ GFP^-^ parent strain. Additional copies of the *ASC1* gene were introduced on a *LEU2* plasmid to avoid recessive mutations in the native *ASC1* gene. To obtain mutants with increased frameshifting efficiency, Ura^+^ mutants were selected and screened for increased GFP/RFP. **(B)** Flow cytometry scatter plot showing GFP versus RFP for 3 mutants and the wild-type parent strain. Expression of *GLN4*_(1-99)_-(CGA)_4_+1-GFP is increased in these *MAT****a*** Ura^+^ mutants. P15: *mbf1-R89K*, P25: *mbf1*Δ*125-151*, P38: *mbf1-K64E*. **(C)** Expression of the non-native tRNA^Arg(UCG)*^ suppressed the Ura^+^ phenotype of mutant P25. Serial dilutions of the indicated strains with empty vector or expressing the mutant tRNA^Arg(UCG)*^ were grown on the indicated media. **(D)** Mutations in the *MBF1* mutants map in conserved amino acids in both the *MBF1*-specific domain and the Helix-Turn-Helix (HTH) domain of Mbf1 protein. Alignment of yeast Mbf1 amino acids 60-100 with other eukaryotic species is shown (full alignment see figure 1-supplement 3A). GFP/RFP of frameshifted (CGA)_4_+1 reporter is shown for mutants obtained from *MAT****a*** (circles) and *MATα* (triangles) strains, with the color of markers corresponding to the consensus level of this residue (Blue: 50%-90%, Red: 90%), however the conserved residue for R61 is N, and for S86 is Q, with all others identical to yeast.

To confirm that inhibitory decoding of CGA codon repeats is required for frameshifting in these mutants, we showed that introduction of an anticodon-mutated exact match tRNA^Arg(UCG)*^ suppressed the Ura^+^ phenotype of one mutant (Fig. 1C). We have shown previously that expression of this exact match tRNA^Arg(UCG)*^ results in efficient decoding of CGA codons and suppresses their inhibitory effects on gene expression (Letzring et al., 2010). Thus, the Ura^+^, GFP^+^ phenotype of this mutant was due to frameshifting that occurs when the ribosome translates CGA codon repeats inefficiently.

We demonstrated that mutations in the yeast gene *MBF1*, Multiprotein-Bridging Factor 1, were responsible for the defects in reading frame maintenance in recessive high GFP mutants. We identified the mutated gene by complementation of the Ura^+^ phenotype of the P25 recessive mutant with two plasmids from a library that contain 97.2% of the entire yeast genome (Fig. 1-figure supplement 1B) (Jones et al., 2008). The complementing plasmids share a single ORF, *MBF1*.

We confirmed that mutations in the *MBF1* gene are responsible for frameshifting in three ways. First, a plasmid with only the *MBF1* gene complemented the frameshifting Ura^+^ phenotype of two mutants (Fig. 1-figure supplement 2A). Second, deletion of *MBF1* in the parent strain converted that strain from GFP^-^ to GFP^+^, similar to deletion of *ASC1* (Fig. 1-figure supplement 2B). Third, 19/19 mutants tested contain mutations in the *MBF1* gene, some of which are shown with frameshifted GFP/RFP values in Figure 1D.

*MBF1* is a highly conserved gene in eukaryotes and archaea (Liu et al., 2007, Takemaru et al., 1997)(Fig. 1-figure supplement 3A), generally <160 amino acids with an N-terminal Mbf1-specific domain and a cro-like helix-turn-helix (HTH) domain (Fig. 1D) (Takemaru et al., 1997). Point mutations isolated in our selection are located at conserved residues near the junctions between the domains (Fig. 1D). Mbf1, which was initially identified as a transcription co-activator in *Bombyx mori* (Li et al., 1994, Takemaru et al., 1997), has a similar function in yeast, in this case, interacting with the Gcn4, transcription regulator of the general amino acid control pathway (Takemaru et al., 1998). In testing sensitivity to 3-aminotriazole (3-AT), a phenotype of *gcn4* mutants due to inability to induce expression of *HIS3*, we found that two frameshifting point mutants (*mbf1-K64E* and *mbf1-I85T*) exhibit no growth defect even on high concentrations of 3-AT (Fig. 1-figure supplement 3B). Moreover, deletion of *GCN4* does not affect frameshifting at CGA codon repeats in an *asc1*Δ mutant (Wolf and Grayhack, 2015). Thus, it is unlikely that the defect in reading frame maintenance in our *mbf1* mutants is related to *GCN4*. However, Mbf1 has also been implicated in translation, based on isolation of mutations in yeast *MBF1* that suppress frameshifting mutations (Hendrick et al., 2001), and the weak association of the archaeal homolog with ribosomes (Blombach et al., 2014). However, there is no information on its molecular role in translation.

### Ribosomal protein *Rps3* also mediates reading frame maintenance at CGA codon repeats

To identify the mutated gene(s) in our dominant mutants, we performed whole genome sequencing in two *MATα* mutants and found that each mutant contains a single amino acid change (*S104Y* and *G121D*) in *RPS3*. Similarly, the two dominant *MAT****a*** mutants also contain mutations in the *RPS3* gene (*L113F* and a duplication of N22 to A30). *RPS3* encodes a universally conserved ribosomal protein, a core component of the mRNA entry tunnel with a eukaryotic-specific C-terminal extension that interacts with Asc1 (Rabl et al., 2011). One mutation in *RPS3* (*K108E*) affects reading frame maintenance (Hendrick et al., 2001), while others affect different aspects of translation, from initiation to quality control (Dong et al., 2017, Graifer et al., 2014, Limoncelli et al., 2017, Takyar et al., 2005). The three residues *S104*, *L113* and *G121* implicated in reading frame maintenance in our study, as well as *K108*, are all found in two α-helices near the mRNA entry tunnel of the ribosome; these residues reside on the surface of the ribosome and could interact with mRNA or proteins outside of the ribosome (Fig. 2A). Furthermore, for all four of these residues, their identity is conserved in eukaryotes and different in bacteria and archaea (Graifer et al., 2014).

**Figure 2.**
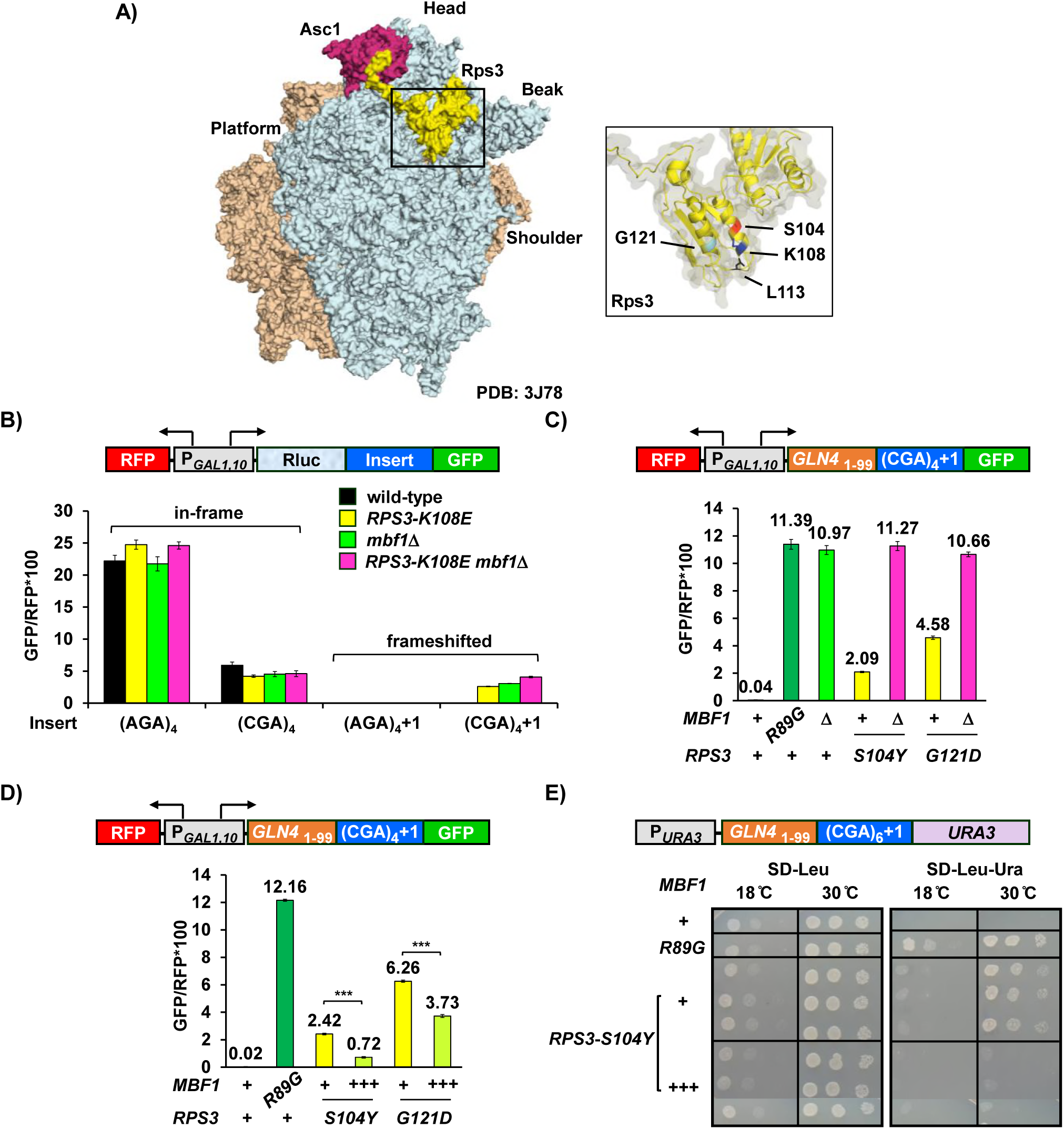
Ribosomal protein Rps3 has a shared function with Mbf1 in preventing frameshifting at CGA codon repeats. **(A) Left:** Yeast ribosome from PDB: 3J78 (Svidritskiy et al., 2014) (light blue: small subunit; sepia: large subunit) showing Asc1/RACK1 (magenta) and Rps3 (yellow). **Right:** Residues of Rps3 in which mutations ca use frameshifting are marked- *S104* (red), *K108* (dark blue), *L113* (black), *G121* (light blue). **(B)** Analysis of effects of *RPS3-K108E, mbf1*Δ *and RPS3-K108E mbf1*Δ mutations on expression of GFP reporters containing four Arg codons (AGA versus CGA) in frame and in the +1 frame. The *K108E* mutation in *RPS3* allows frameshifting CGA codon repeats, and the combined effects of *RPS3*-*K108E* and *mbf1*Δ are not additive. **(C)** Epistatic assay of *RPS3* mutations from this selection and the *mbf1*Δ strain indicated that these *RPS3* mutations allow frameshifting at CGA codon repeats and do not increase frameshifting in *mbf1*Δ mutants. **(D)** Overproduction of Mbf1 protein in indicated *RPS3* mutants significantly decreased expression of frameshifted Gln4-GFP fusion protein (*** p<0.001) analyzed by flow cytometry. **(E)** Overproduction of Mbf1 protein in the *RPS3-S104Y* mutant reduced frameshifting-dependent growth on -Ura media, shown by a spot test assay.

We initially examined the effect of the *RPS3-K108E* mutation on frameshifting and read-through at CGA codon repeats, and found that this mutation does allow frameshifting but does not affect read-through. To this end, we introduced modified RNA-ID reporters into *rps3*Δ::*ble^R^* strains in which the only source of *RPS3* is a plasmid borne copy (either wild type or *K108E*). As described previously, since the expression of GFP and RFP are driven by the bi-directional *GAL1, 10* promoter, we use the ratio of GFP/RFP to reduce noise and cell type specific differences in induction of this promoter (Dean and Grayhack, 2012). We found that neither the *RPS3* mutant nor an *mbf1*Δ mutant affected in frame read-through of CGA codon repeats (Fig. 2B). However, both the *RPS3*-*K108E* and *mbf1*Δ mutants caused increased expression of frameshifted GFP in the construct with four CGA codons (Fig. 2B; Supplementary Table 1). Since the *K108E* mutation has only minor effects on the polysome to monosome ratio (Dong et al., 2017), we infer that effects of this mutation are specific to reading frame maintenance (Fig.2B).

If Mbf1 and Rps3 proteins function in independent pathways to prevent frameshifting, we expected that *RPS3-K108E mbf1*Δ double mutants would frameshift more efficiently than either single mutant. Instead, we found that the double mutant *RPS3-K108E mbf1*Δ exhibited only a slight increase in expression of frameshifted GFP, much less than an additive effect (Fig. 2B). We also compared expression of *GLN4*_(1-99)_-(CGA)_4_+1-GFP in the *MATα mbf1-R89G* mutant, two *MATα RPS3* mutants (*S104Y* and *G121D*) from our selection, in an *mbf1*Δ mutant, and in each *RPS3 mbf1*Δ double mutant. We observed significant amounts of frameshifted GFP in both *RPS3* mutant strains and in the *mbf1*Δ strain as well as in the *mbf1*-*R89G* mutant (Fig. 2C). In these cases again, the double mutants exhibited similar amounts of frameshifted GFP/RFP, compared to the *mbf1*Δ strain, although an additive effect would be easily detectable (Fig. 2C). Thus, we think it is likely that Mbf1 and the two α-helices in the N-terminal Rps3 protein have related roles in reading frame maintenance.

If Mbf1 and these two α-helices in Rps3 mediate a common function, then frameshifting in either *RPS3*-*S104Y* or *G121D* mutants might be suppressed by overproduction of *MBF1*. We find that introduction of additional copies of the *MBF1* gene into either of these mutants resulted in reduced expression of frameshifted GFP (Fig. 2D). Frameshifted GFP is reduced to 30% in the *S104Y* mutant and to 60% in the *G121D* mutant (Fig. 2D). Similarly, growth on media lacking uracil is severely compromised in the *RPS3-S104Y* mutant when *MBF1* is expressed on a multi-copy plasmid, relative to an empty vector control (Fig. 2E), although both strains grow equally well on SD-Leu media. These observations are consistent with the idea that Mbf1 and Rps3 play similar roles in reading frame maintenance and support the idea that these *RPS3* mutations reduce Mbf1 function.

### Mbf1 and Asc1 play distinct roles at CGA codon repeats

Since Asc1 is also required for reading frame maintenance at CGA codon repeats (Wolf and Grayhack, 2015), we examined the relationship between *MBF1* and *ASC1 by* comparing the frameshifting efficiency as well as in-frame read-through in the *asc1*Δ *mbf1*Δ double mutant to that in either single mutant. Since we previously noted that inhibitory effects of CGA codons are mediated by CGA codon pairs (Gamble et al., 2016, Letzring et al., 2010), we compared effects of these mutants on a set of reporters with three CGA-CGA (or AGA-AGA) codon pairs flanked by two non-Arg codons (Fig. 3A; Supplementary Table 2) to effects on a set with four adjacent CGA (or AGA) codons (Fig. 3-figure supplement 1).

**Figure 3.**
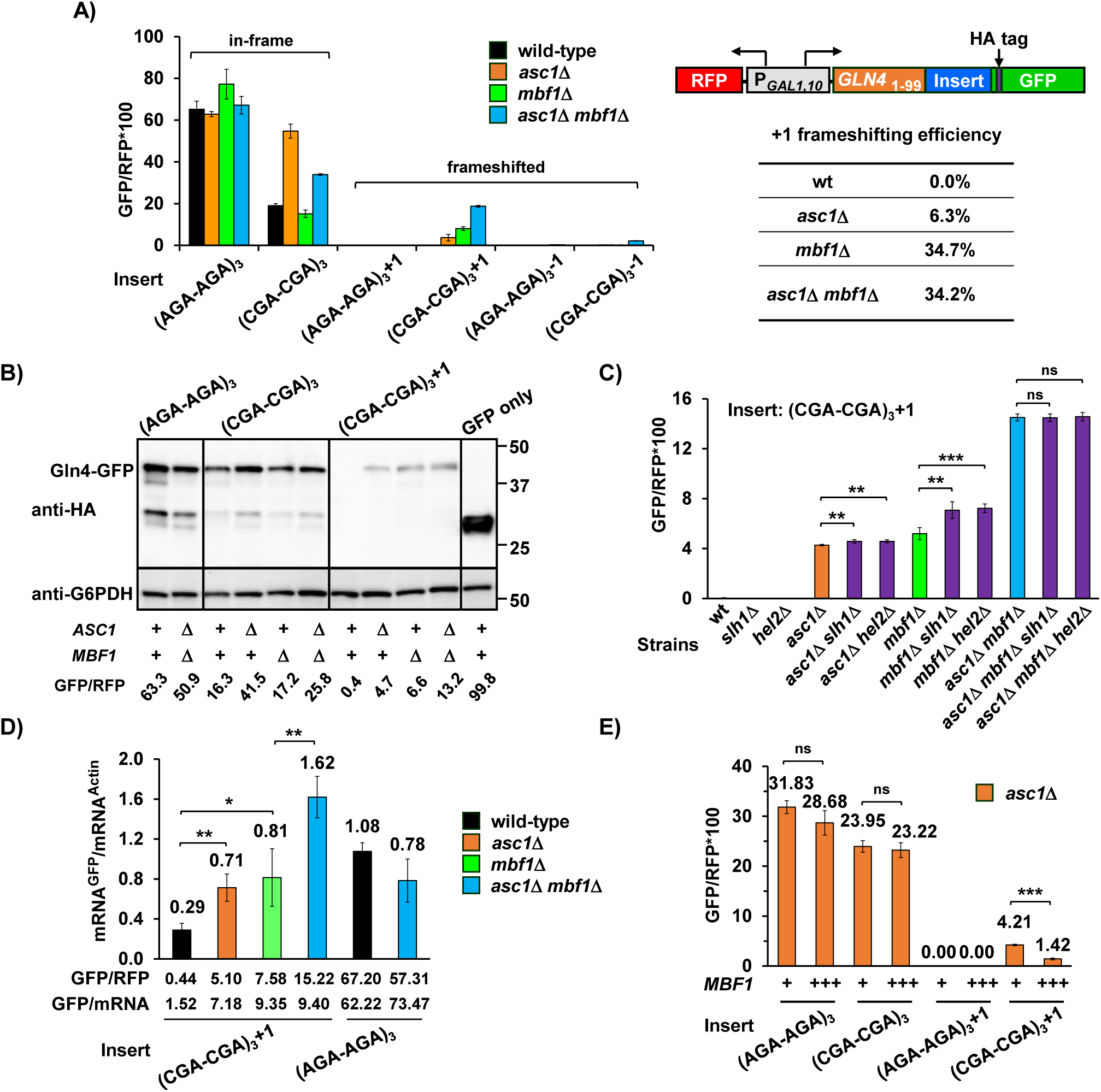
Mbf1 and Asc1 play distinct roles at CGA codon pairs. **(A)** Analysis of effects of *asc1*Δ*, mbf1*Δ *and asc1*Δ *mbf1*Δ mutations on expression of *GLN4*_(1-99)_-GFP reporters containing three Arg-Arg codon pairs (AGA-AGA versus CGA-CGA) in 0, +1, and −1 reading frames. Mutation of either *ASC1* or *MBF1* allows frameshifting in the (CGA-CGA)_3_+1 reporter, and mutation of both *ASC1* and *MBF1* results in significantly more frameshifted GFP/RFP. The +1 frameshifting efficiency [(+1 GFP/RFP) / (+1 GFP/RFP + in-frame GFP/RFP + −1 GFP/RFP)] of all four strains is shown in the table. **(B)** Western analysis of Gln4-GFP fusion protein in yeast strains from (A) indicates the expression of frameshifted Gln4-GFP full-length protein in all three mutants. The protein was detected by anti-HA antibody recognizing the HA epitope between the codon insert and GFP. The GFP and RFP values were measured by flow cytometry while harvesting for cell lysis. **(C)** Effects of *hel2*Δ *and slh1*Δ on frameshifting at CGA-CGA codon pairs with and without deletions in *MBF1* and *ASC1*. ns: p>0.05, * p<0.05, ** p<0.01, *** p<0.001 **(D)** Analysis of the mRNA levels of the *GLN4*-GFP reporter by RT-qPCR. Deletion of *ASC1* and/or *MBF1* resulted in increased mRNA. * p<0.05, ** p<0.01 **(E)** Overproduction of Mbf1 suppressed frameshifting at CGA-CGA codon pairs in the *asc1*Δ mutant, but did not affect the in-frame read-through, based on GFP/RFP expression from the indicated reporters shown in (A). ns: p>0.05, *** p<0.001.

We found that Asc1, but not Mbf1, mediates the inhibition of translation conferred by CGA-CGA codon pairs, and that neither the upstream gene nor the arrangement of CGA codons affected this result. While deletion of *ASC1* resulted in increased in-frame expression of both CGA-containing reporters, deletion of *MBF1* did not, in fact a small decrease in GFP/RFP is observed (Fig. 3A; Fig. 3-figure supplement 1; Supplementary Table 2). The double deletion strain exhibited an intermediate level of in-frame GFP expression (Fig. 3A).

If Mbf1 and Asc1 proteins function in independent pathways that affect frameshifting at CGA codon pairs, we expected that *asc1*Δ *mbf1*Δ double mutants would frameshift more efficiently than either single mutant. Frameshifting occurs in both the single and double mutants (*asc1*Δ, *mbf1*Δ, *asc1*Δ *mbf1*Δ), but the amount of frameshifted GFP protein in the double mutant was greater than the sum of frameshifted GFP produced in two single mutants (Fig. 3A; Fig. 3-figure supplement 1; Supplementary Table 2). Moreover, in the double mutant, a small amount of frameshifting is also detected in the −1 frame (Fig. 3A; Fig. 3-figure supplement 1; Supplementary Table 2). We confirmed that +1 GFP signal detected in our mutants was due to frameshifting rather than another aberrant translation event by directly measuring both the size and amount of GFP fusion protein. The amount of full-length GFP protein in the Western blot corresponds to the GFP/RFP values obtained from flow analysis (Fig.3B) indicating that +1 GFP/RFP signal in our mutants is due to frameshifting.

We have three results that are consistent with an important role for Asc1 in the decision between read-through versus activation of the RQC pathway, rather than a major direct role in reading frame maintenance. First, the deletion of *ASC1* in an *mbf1*Δ mutant does not affect the frameshifting efficiency of the (CGA-CGA)_3_ constructs, but rather affects the total number of ribosomes translating through the CGA codons. We calculate frameshifting efficiency as the percentage of GFP/RFP from the +1 construct relative to the total GFP/RFP from the in frame, +1 and −1 constructs with the same insert [(+1 GFP/RFP) *100 / (+1 GFP/RFP + in-frame GFP/RFP + −1 GFP/RFP)]. For the *GLN4*_(1-99)_-(CGA-CGA)_3_-GFP reporter, 34% of the GFP signal corresponds to the +1 frameshift in both *mbf1*Δ and *asc1*Δ *mbf1*Δ mutants (Fig. 3A, Supplementary Table 2), although this is not true for the *Renilla* luciferase-(CGA)_4_-GFP reporters perhaps due to a previously observed effect of Asc1 on *Renilla* luciferase (Fig. 3-figure supplement 1). Second, other mutations that impair the recruitment of the RQC pathway, but do not themselves affect frameshifting, also result in increased amount of frameshifted GFP in *mbf1*Δ strains. Frameshifting was increased by deletions of either of two downstream effectors of Asc1 (*HEL2* or *SLH1)* in an *mbf1* mutant, although neither of these single mutants allows detectable frameshifting (Fig. 3C; Supplementary Table 3) (Wolf and Grayhack, 2015), while both single mutants increase read-through (Brandman et al., 2012, Sitron et al., 2017). Third, the amount of frameshifted GFP per mRNA is constant between *mbf1*Δ versus *asc1*Δ *mbf1*Δ mutants. Frameshifting was proportional to the abundance of the *GLN4*_(1-99)_-(CGA-CGA)_3_+1-GFP mRNA in *mbf1*Δ versus *asc1*Δ *mbf1*Δ mutants (Fig. 3D), although the mRNA in the *asc1*Δ *mbf1*Δ mutant was twice that in the *mbf1*Δ single mutant (Fig. 3D). Thus, we infer that Asc1 mediates the balance between read-through and aborted translation at CGA codon repeats, and that aborted translation helps to maintain the reading frame. These results are consistent with an important role for Asc1 in the decision between read-through versus activation of the RQC pathway, while Mbf1 acts solely on reading frame maintenance.

If Mbf1 is responsible for reading frame maintenance in all conditions, then overproduction of Mbf1 in the *asc1*Δ mutant might suppress frameshifting in this mutant. We find that expression of *MBF1* on a multi-copy plasmid did suppress frameshifting in the *asc1*Δ strain to 1/3 that seen with an empty vector, but did not affect the in-frame read-through (Fig. 3E). The overproduction of Mbf1 is not complementing a reduced abundance of Mbf1 in this mutant. We did not detect a reduction in Mbf1-HA (which complements the *mbf1*Δ mutant) in the *asc1*Δ strain (Fig. 3-figure supplement 2A), although *asc1* mutants generally exhibit a defect in expression of small proteins (Thompson et al., 2016). We also considered that *mbf1* mutants might require additional Asc1 protein, but additional copies of *ASC1* did not suppress frameshifting in an *mbf1*Δ mutant (Fig. 3-figure supplement 2B). We infer that Mbf1 and Asc1 contribute in distinct ways to reading frame maintenance, although we have not ruled out an additional role for Asc1 in reading frame maintenance.

### Mbf1 regulates frameshifting at slowly translated codon pairs, mainly those targeted by Asc1

Most efficient frameshifting occurs at sequences that are slowly translated (Caliskan et al., 2014). We considered that Mbf1 and/or Asc1 might be important for reading frame maintenance at some of the 17 inhibitory codon pairs in yeast that cause reduced expression and exhibit high ribosome occupancy, indicative of slow translation (Gamble et al., 2016). Thus, we examined frameshifting at 12 of 17 inhibitory codon pairs, including 11 of the 12 most slowly translated pairs (Gamble et al., 2016).

We found that ribosomes frameshift at 7 of the 12 inhibitory pairs in the *asc1*Δ *mbf1*Δ double mutant, with high levels of frameshifting at 3 codon pairs (CGA-CGA, CGA-CGG, and CGA-CCG) and low, but distinct, levels at 4 other pairs (CGA-AUA, CGA-CUG, CUC-CCG, and CGA-GCG) (Fig. 4A; Supplementary Table 4). These pairs are the seven most slowly translated codon pairs in the yeast genome, and the only inhibitory or slow pairs with CGA in the 5’ position (Gamble et al., 2016). We noted that in the *mbf1*Δ strain, significant frameshifting occurs only at the 3 pairs with the highest frameshifting levels in the *asc1*Δ *mbf1*Δ double mutant.

**Figure 4.**
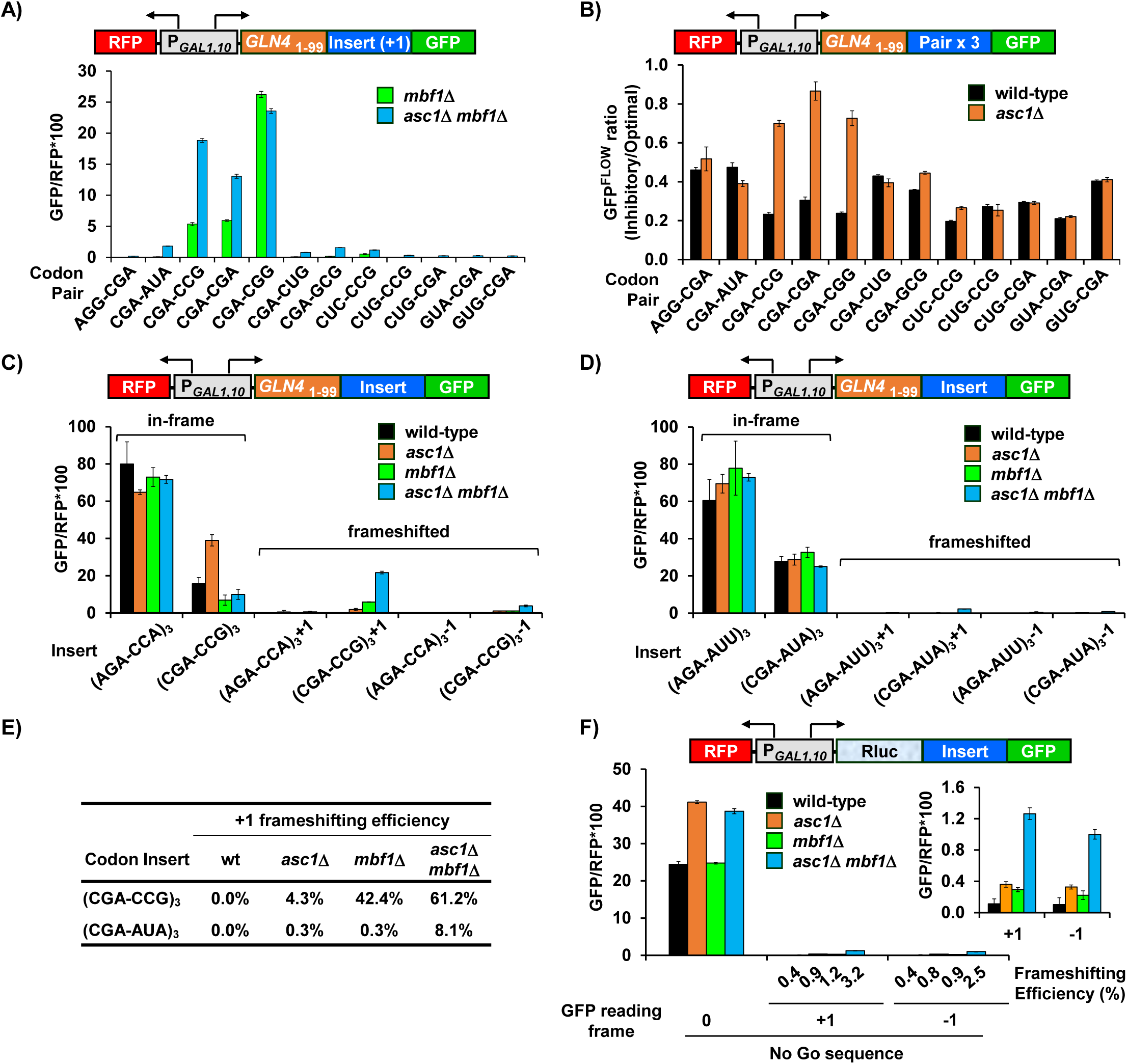
Mbf1 regulates frameshifting a t slowly translated inhibitory codon pairs, mainly those targeted by Asc1. **(A)** Frameshifting is detected at three inhibitory codon pairs (Gamble et al., 2016) in the *mbf1*Δ mutant, and at seven codon pairs in the *asc1*Δ *mbf1*Δ double mutant. Frameshifting was assayed from reporters bearing 3 copies of the indicated inhibitory codon pair and a +1 nucleotide to place GFP in the +1 frame. **(B)** In frame read1064 through of three inhibitory codon pairs (CGA-CGA; CGA-CCG; CGA-CGG) is improved by the deletion of *ASC1*. GFP/RFP from reporters with three copies of an inhibitory pair were compared to synonymous reporters with three copies of the optimized pair to obtain GFP^FLOW^ ratio. **(C**, **D)** Analysis of effects of *asc1*Δ*, mbf1*Δ *and asc1*Δ *mbf1*Δ mutations on expression of *GLN4*-GFP reporters containing three copies of either (C) the Arg-Pro (AGA-CCA or CGA-CCG) codon pairs or (D) the Arg-Ile (AGA-AUU or CGA-AUA) codon pairs in 0, +1, and −1 reading frames. Mutation of either *ASC1* or *MBF1* allows frameshifting in the (CGA-CCG)_3_+1 reporter, but not in the (CGA-AUA)_3_+1 reporter, while mutations of both *ASC1* and *MBF1* results in significantly more frameshifted GFP/RFP in both reporters. **(E)** The +1 frameshifting efficiency at either CGA-CCG codon pairs or CGA-AUA codon pairs in all four strains is shown. **(F)** Mutation of either *ASC1* or *MBF1* allows frameshifting at No-Go sequences in the GFP reporter, and mutation of both *ASC1* and *MBF1* results in significantly more frameshifted GFP/RFP.

Since Asc1 primarily regulates read-through of CGA codon pairs, we considered that Asc1 might have a similar role at other inhibitory codon pairs, explaining why high levels of frameshifting occur in the *asc1*Δ *mbf1*Δ double mutant. We found that deletion of *ASC1* resulted in increased in-frame read-through of 3 inhibitory pairs (CGA-CGA, CGA-CGG, and CGA-CCG) (Fig. 4B; Supplementary Table 4), the 3 pairs that exhibited high levels of frameshifting in both the *asc1*Δ *mbf1*Δ double mutant and the *mbf1*Δ strain. In each case, we measured the expression of each inhibitory codon pair relative to its synonymous optimal pair, obtaining a GFP^FLOW^ ratio (Gamble et al., 2016) in wild type and *asc1*Δ mutants. Therefore, Asc1 mediates inhibitory effects of only a subset of the slowly translated inhibitory codon pairs. Moreover, deletion of *ASC1* is important for frameshifting at four pairs for which Asc1 has little (CGA-GCG; CUC-CCG) or no effect on read-through. The basis for Asc1 regulation of particular codon pairs is unknown, since the dependence upon Asc1 does not correlate strictly with cumulative ribosome occupancy, the A-P ribosome occupancy (Gamble et al., 2016) or the ratio of long to short footprints described by Matsuo *et al*. (Matsuo et al., 2017).

We infer that Asc1 may have a role in frameshifting, in addition to its effects on read344 through, based on examination of frameshifting and in-frame read-through at CGA-CCG and CGA-AUA pairs (Fig. 4C, 4D, 4E; Supplementary Table 2). For the CGA-CCG pair, we detected significant +1 frameshifting with the CGA-CCG pair in both the *asc1*Δ and *mbf1*Δ single mutants, but the +1 GFP in the *asc1*Δ *mbf1*Δ mutant was more than double the sum of the +1 GFP in each single mutant (Fig. 4C); frameshifting efficiency increased from 42.4% in the *mbf1*Δ to 61.2% in the *asc1*Δ *mbf1*Δ mutant (Fig. 4E). Even more surprisingly, for the CGA-AUA pair, frameshifting efficiency increased from 0.3% in the *mbf1*Δ and *asc1*Δ single mutants to 8.1% in the *asc1*Δ *mbf1*Δ (Fig. 4D, 4E). Thus, Asc1 may have an additional role in frameshifting that is not a simple extension of its role in aborting translation.

### Mbf1 regulates frameshifting at other slowly translated sequences

Since ribosomes frameshift at the 7 most slowly-translated inhibitory codon pairs in the *asc1*Δ *mbf1*Δ double mutant, we hypothesized that any slowly translated sequence might provoke frameshifting in this mutant. To test our hypothesis, we measured frameshifting at a sequence which forms secondary structure to slow down translation and induce No-Go mRNA decay (Doma and Parker, 2006, Harigaya and Parker, 2010, Passos et al., 2009). We found that frameshifting occurred in both directions, and was detectable in wild type, greater in each single mutant and even greater in the *asc1*Δ *mbf1*Δ mutant (Fig. 4F; Supplementary Table 5). By contrast, we did not observe an increase in frameshifting efficiency at the programmed frameshift site for *TY1* (Fig. 4-figure supplement 1; Supplementary Table 5), in which a translational pause at an Arg AGG codon decoded by a rare tRNA allows slippage between a Leu CUU codon (in frame) and a UUA codon (in the +1 frame) (Belcourt and Farabaugh, 1990). Thus, Mbf1 regulates reading frame maintenance at a translational pause (No-Go site), but does not enhance frameshifting at site in which translational slippage is encoded.

### Efficient frameshifting occurs at single CGA-CGG pair in a particular context

Frameshifting at the CGA-CGG codon pair yielded the most frameshifted +1 GFP and exhibited similar amounts of +1 GFP in the *mbf1*Δ mutant and in the *asc1*Δ *mbf1*Δ mutant (Fig. 4A). Thus, we examined expression of a complete set of reporters and found that frameshifting efficiency with these CGA-CGG constructs was ~35% in the *asc1*Δ, ~76% in the *mbf1*Δ and ~55.4% in the *asc1*Δ *mbf1*Δ double mutants (Fig. 5-figure supplement 1A; Supplementary Table 2). These results were somewhat surprising since CGA-CGG is neither as inhibitory nor as slowly translated as either the CGA-CCG or CGA-CGA pair.

**Figure 5.**
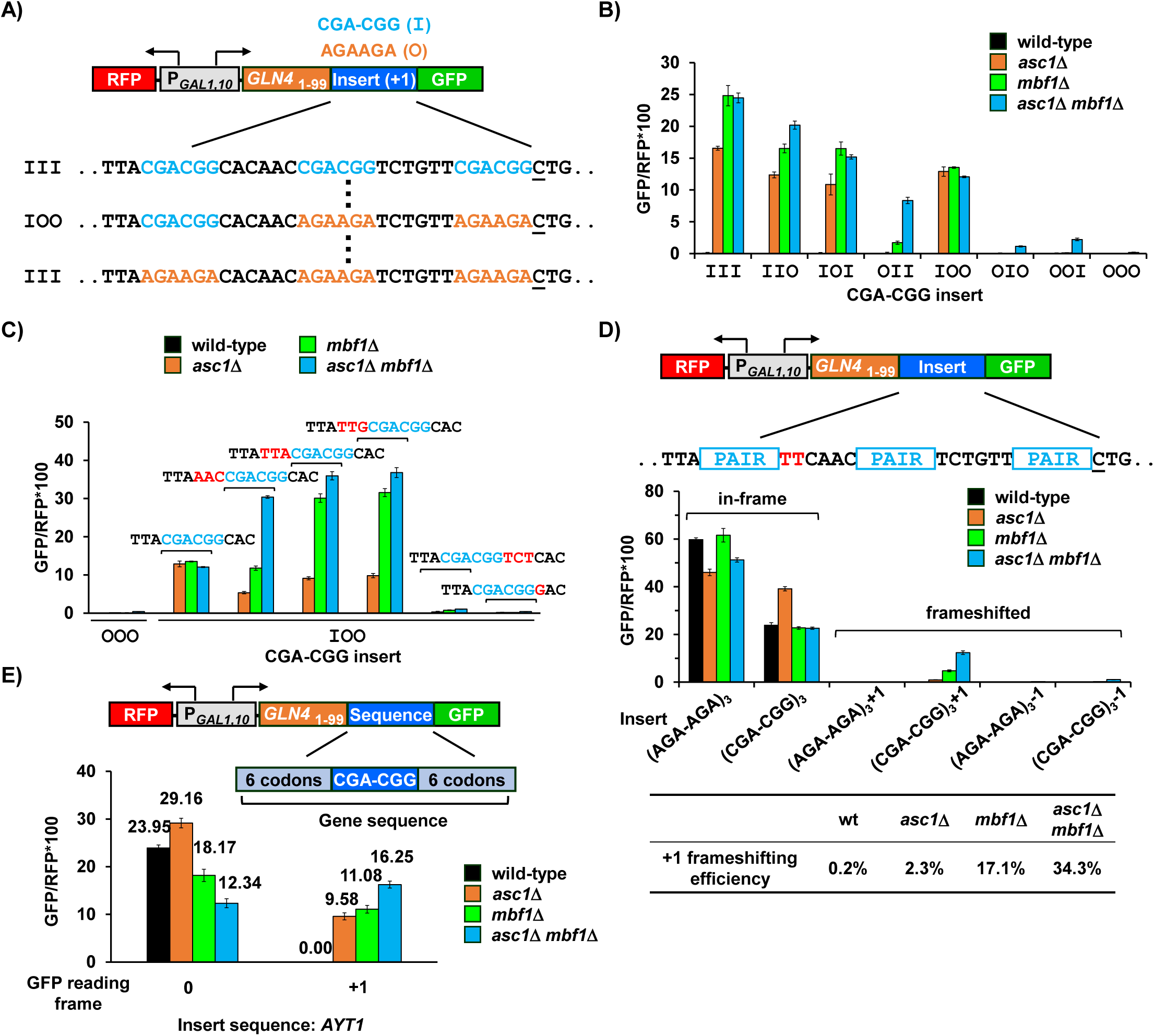
Efficient frameshifting occurs at a single CGA-CGG pair in a particular context. **(A)** Schematic of inserts in modified RNA-ID reporters used to identify the contributions of individual CGA-CGG pairs to frameshifting. Sequences with all possible combinations of zero, one, two or three inhibitory CGA-CGG pairs (I, shown in cyan) [substituting the synonymous optimal pair AGA-AGA (O, shown in orange) at other positions] were inserted between *GLN4*_(1-99)_ and GFP. **(B)** Analysis of effects of *asc1*Δ*, mbf1*Δ *and asc1*Δ *mbf1*Δ mutations on expression of *GLN4*_(1-99)_-GFP reporters with the indicated position and number of inhibitory codon pairs. All constructs with an inhibitory codon pair at the first position (III, IIO, IOI, IOO) showed high levels of frameshifting in all three mutants. **(C)** Analysis of *GLN4*_(1-99)_-GFP reporters with IOO CGA-CGG construct in which the sequences surrounding the single CGA-CGG insert were varied. The 3’ nucleotide of the first CGA-CGG pair is required for efficient frameshifting in the mutants. All changes are shown in red. **(D)** Analysis of effects of *asc1*Δ*, mbf1*Δ *and asc1*Δ *mbf1*Δ mutations on expression of revised *GLN4*_(1-99)_-GFP reporters (TTC is substituted for CAC as the 3’ codon downstream of the first codon pair) containing three Arg-Arg codon pairs (AGA-AGA versus CGA-CGG) in 0, +1, and −1 reading frames. Mutation of either *ASC1* or *MBF1* allows frameshifting in this (CGA-CGG)_3_+1 reporter, and mutation of both *ASC1* and *MBF1* results in significantly more frameshifted GFP/RFP. The +1 frameshifting efficiency of all four strains is shown in the table. **(E)** Analysis of effects of *asc1*Δ*, mbf1*Δ *and asc1*Δ *mbf1*Δ mutations on expression of *GLN4*_(1-99)_-GFP reporters containing the native yeast *AYT1* sequence with a single CGA-CGG codon pair in 0 and +1 reading frames. This native yeast sequence provoked significant amount of frameshifting in the *asc1*Δ *mbf1*Δ strain with small reduction of in-frame read-through.

We defined the contributions to frameshifting of each CGA-CGG codon pair in the three codon pair construct, because this analysis might point to particular contexts that affect frameshifting efficiency. Moreover, the reduced frameshifting associated with fewer inhibitory codon pairs might restore synergistic effects of *ASC1* and *MBF1*. To this end, we constructed reporters with all possible combinations of zero, one, two, or three CGA-CGG (I) pairs [substituting the synonymous optimal pair AGA-AGA (O) at other positions] (Fig. 5A). We found that all constructs with an inhibitory codon pair at the first position (III, IIO, IOI, IOO) showed high levels of frameshifting in all three mutants and little synergy of the double mutant (Fig. 5B; Supplementary Table 6). By contrast, constructs with an optimal codon pair at the first position (OII, OIO, OOI) showed low levels of frameshifting in either single mutant and enhanced frameshifting in the double mutant (Fig. 5B), consistent with results with other pairs. Thus, we infer that combination of CGA-CGG and the particular sequence context of the first position is responsible for highly efficient frameshifting.

To discern the requirements for efficient frameshifting, we analyzed a set of variants of the CGA-CGG IOO construct altering a codon or nucleotide upstream or downstream of the CGA-CGG pair. We found that the CGA-CGG-C 7-mer is required for efficient frameshifting. Either of two changes to the sequences downstream of the CGA-CGG pair (one a point mutation and another a codon insertion) eliminated efficient frameshifting in all three mutant strains (Fig. 5C). By contrast, insertions of any of three codons upstream of the CGA-CGG pair did not eliminate efficient frameshifting in the *mbf1*Δ or *asc1*Δ *mbf1*Δ mutants (Fig. 5C; Supplementary Table 7). In fact, all upstream changes enhanced frameshifting in the *asc1*Δ *mbf1*Δ mutant, two did so in the *mbf1*Δ mutant, while all three changes reduced frameshifting in the *asc1*Δ mutant. These observations suggest that there are differences in the requirements for frameshifting in different mutants. Thus, we infer that the CGA-CGG-C 7-mer is required for efficient frameshifting, but we note that CGA-CGG-C 7-mer is not sufficient for efficient frameshifting since the third CGA-CGG is also followed by a C. Furthermore, we restored the synergistic interaction between *MBF1* and *ASC1* by simply altering the downstream nucleotides from CA to TT in a three codon pair reporter (Fig. 5D; Supplementary Table 2).

We investigated frameshifting at native gene sequences that contain CGA-CGG codon pairs to find out if *mbf1*Δ, or *asc1*Δ *mbf1*Δ mutants allowed frameshifting in this context. Expression of frameshifted fusion protein was detectable with sequences from 7/7 tested genes in the double mutant, with frameshifted GFP/RFP ranging from 0.24 to 16.3 (Fig. 5E; Fig. 5-figure supplement 1B; Supplementary Table 8). The levels of frameshifted GFP do not correlate with the levels of in-frame expression, since the highest levels of frameshifted GFP were observed with the *AYT1* gene, in which in-frame GFP/RFP (9.8) in the *asc1*Δ *mbf1*Δ mutant was actually less than frameshifted GFP/RFP. *AYT1* is one of two genes with CGA-CGG-C sequence and the sequence GTA-CGA-CGG contains two adjacent inhibitory codon pairs. Thus, relatively small native sequences suffice to promote frameshifting at different levels.

### +1 frameshifting occurs with the CGA codon in the P site

To understand how frameshifting occurs, we wanted to define the direction and position of the actual frameshift. The high efficiency of frameshifting at the CGA-CGG-CAC sequence provided a useful tool to study frameshifting since there is only a short potential frameshifting sequence (a single inhibitory codon pair). We inserted this sequence with its neighboring codons from the RNA-ID reporter into a construct for purification of the frameshifted polypeptide (Fig. 6A). The construct was designed such that the protein could be purified either with an upstream affinity tag (GST) to yield all polypeptides or with a downstream affinity tag (Strep II; ZZ domain of IgG) to yield only frameshifted polypeptides. Treatment with LysC, which cleaves after lysine was expected to yield a 16-17 amino acid peptide for analysis by mass spectrometry.

**Figure 6.**
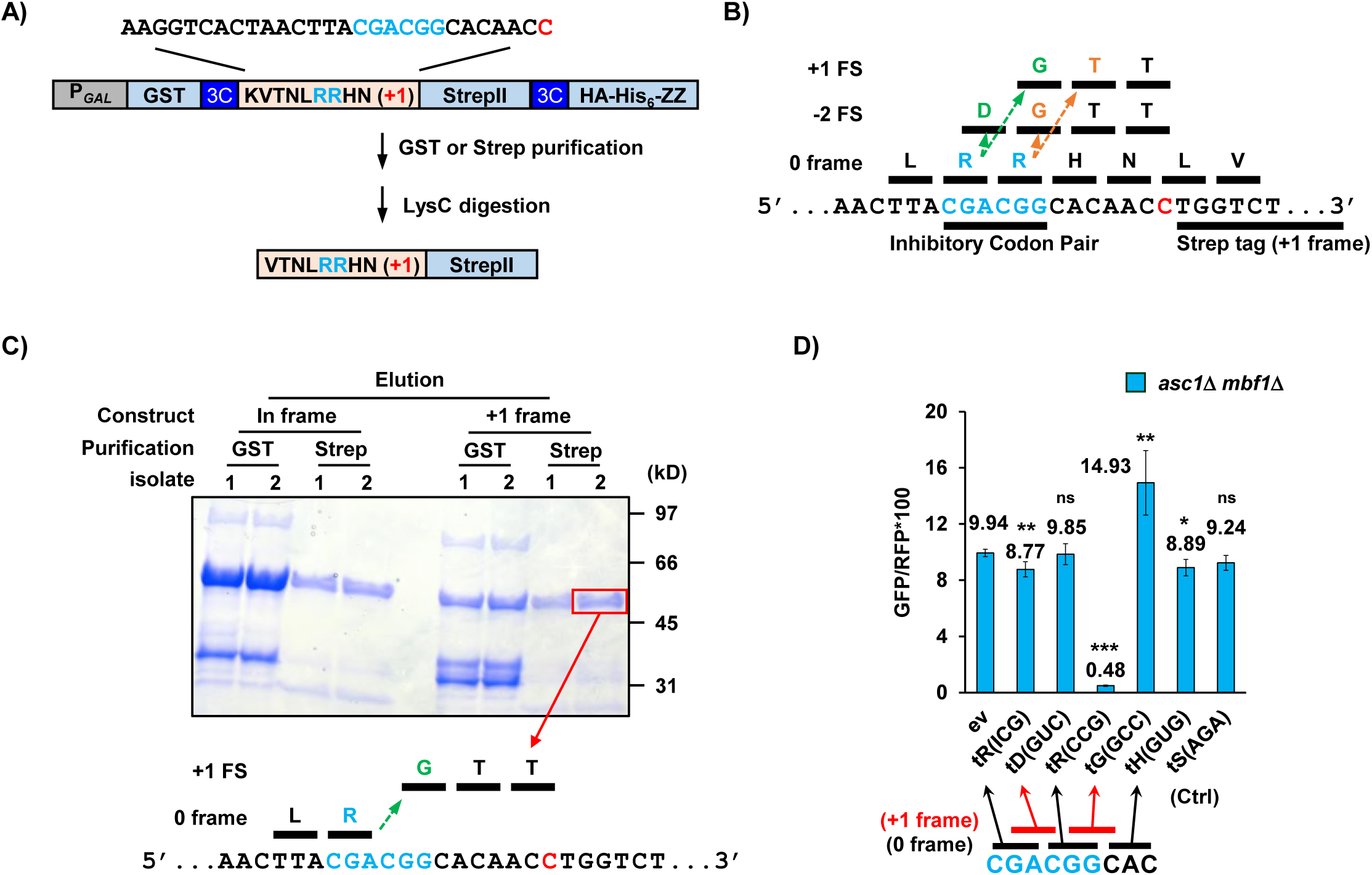
Frameshifting occurs in the +1 direction with the CGA codon in the P site and is modulated by tRNA competition at the A site. **(A)** Schematic of purification construct for frameshifted peptide. An eight amino acid sequence with a single CGA-CGG pair from the RNA-ID reporter was inserted between a GST tag and an out-of-frame StrepII tag. LysC treatment of purified frameshifted protein yields a 16 or 17 amino acid peptide. The red nucleotide indicates the extra nucleotide in the +1 frame construct. **(B)** Schematic of four possible frameshifting events at the inhibitory CGA-CGG codon pair, each of which can be distinguished by one or two amino acids in the resulting peptide. Ribosomes can frameshift either in the forward direction (+1) or in the reverse direction (−2) when the P site is occupied by either the CGA codon (first amino acid in the out-of-frame peptide shown in green) or the CGG codon (first amino acid of out-of-frame peptide shown in orange). **(C)** Purified protein products of both in frame and +1 frame constructs were analyzed by SDS-PAGE, stained with Coomassie Blue. The frameshifted protein of +1 frame construct from Strep purification (in red box) was excised, cleaved with LysC and analyzed by Mass Spectrometry, resulting in identification of the peptide shown below the figure. This peptide corresponds to that expected of a +1 frameshift occurring when the CGA codon occupies the P site. **(D)** Overexpression of tRNA corresponding to +1 frame codon improved frameshifting efficiency, while overexpression of tRNA corresponding to next in frame codon significantly reduced frameshifting. ns: p>0.05, * p<0.05, ** p<0.01, *** p<0.001.

If frameshifting occurred in the local region near the CGA-CGG codon pair, there are four possible events that could all give rise to +1 GFP signal. Ribosomes could frameshift in the +1 direction with either the CGA or the CGG in the P site, yielding the RGTT or the RRTT sequences shown in Fig. 6B. Alternatively, ribosomes could undergo −2 frameshifting at the either codon, yielding the peptides RDGTT or RRGTT (Fig. 6B). In yeast, −2 frameshifting was observed upon expression of the mammalian antizyme (Matsufuji et al., 1996) and −2 frameshifting also occurs in PRRSV virus (Fang et al., 2012). We purified the frameshifted protein, as well as an in-frame control protein with the sequence expected for a −2 frameshift at CGG (Fig. 6C) and subjected them to mass spectrometry. The frameshifted protein yielded the peptide VTNLRGTTWSHPQFEK, the expected peptide from a +1 frameshift beginning with the CGA codon in the P site of the ribosome. Thus, we infer that frameshift occurs with CGA in the P site, yielding only one Arg amino acid on the nascent peptide, then switches to a glycine codon GGC.

To determine if aminoacyl tRNA amounts affect frameshifting, we compared the effects of additional copies of specific Arg and Gly tRNAs on frameshifting in the *asc1*Δ *mbf1*Δ double mutant. We found that introduction of additional copies of the gene encoding tRNA^Arg(CCG)^, which decoded the in-frame CGG codon, severely reduced frameshifting (Fig. 6D), as expected if arg-tRNA^ArgCCG)^ competes with gly-tRNA^Gly(GCC)^ for the A site. Similarly, we found that addition of extra copies of tRNA^Gly(GCC)^ which decodes +1 frame GGC codon significantly increased frameshifting in our original CGA-CGG-CAC context, as might be expect if the GGC codon is used (Fig. 6D). Additional copies of tRNA^Arg(ICG)^, tRNA^Asp(GUC)^, tRNA^His(GUG)^, tRNA^Ser(AGA)^ had little or no effect, as expected since none of the codons decoded by these tRNAs should be occupying the A site during frameshifting. These results indicate that the frameshifting occurs within the single CGA-CGG-CAC sequence and is modulated by the concentration of aminoacyl tRNAs decoding the out-of-frame codon.

## DISCUSSION

We have uncovered a eukaryotic specific system that inhibits frameshifting by stalled ribosomes, in which reading frame maintenance is achieved in two ways, both by direct inhibition of frameshifting and by aborted translation of stalled ribosomes. The system is composed of two proteins that lack bacterial homologs, the archaeal/eukaryotic Mbf1 protein and the eukaryotic ribosomal protein Asc1/RACK1, as well as one universally conserved ribosomal protein Rps3. In wild type cells, ribosomes stall either due to inhibitory codon pairs or structures within the RNA. Mbf1 and Rps3 cooperate at these stalled ribosomes to prevent frameshifting, which, in turn, allows Asc1 to trigger a set of responses that result in aborted translation and recruitment of the RQC complex. In the absence of Mbf1 and Asc1, ribosomes frameshift efficiently, even at a single CGA-CGG pair in some cases, including sequences found in the native yeast genome. Frameshifting on the CGA-CGG codon pair occurs in the +1 direction, with the CGA codon in the P site of the ribosome and is modulated by availability of in-frame and +1 frame A site tRNAs.

We provide evidence that when ribosomes slow down during translation elongation, two distinct sets of events occur. Mbf1 and Rps3 actively prevent frameshifting, while Asc1 recruits Hel2 and Slh1 to abort translation and recycle the ribosome. These two pathways, a reading frame maintenance system and the RQC pathway, cooperate to keep ribosomes on track. We document four observations that support aspects of this model. First, we find that slow or paused ribosomes require the Asc1 and Mbf1/Rps3 intervention, since frameshifting was observed in the *asc1*Δ *mbf1*Δ double mutant at the seven most slowly translated codon pairs in yeast (all inhibitory codon pairs) and at a sequence known to provoke No-Go decay. Second, we demonstrate that Asc1 and Mbf1 have at least one distinct role with respect to the stalled ribosomes. Only Asc1, but not Mbf1, affects the in-frame read-through at inhibitory codon pairs. Asc1 mediates key processes at the stalled ribosome, including recruitment of Hel2 and Slh1, which in turn recruit the RQC complex (Brandman and Hegde, 2016, Brandman et al., 2012, Joazeiro, 2017, Juszkiewicz and Hegde, 2017, Matsuo et al., 2017, Simms et al., 2017, Sundaramoorthy et al., 2017, Shen et al., 2015, Sitron et al., 2017). Third, Mbf1 and Rps3 work together, based on the observations that the double mutant has little increase in frameshifting relative to either single mutant; overproduction of Mbf1 suppressed frameshifting in two *RPS3* mutants; and mutations in either gene only affect frameshifting, not read-through. Fourth, the Asc1 and Mbf1 pathways each act to prevent frameshifting, because *asc1*Δ *mbf1*Δ double mutants display significantly more frameshifting than either single mutant. Asc1 activity is critical to prevent frameshifting, because ribosomes that do not abort translation through Asc1 action likely remain stalled and have an increased chance of frameshifting.

We think Rps3 and Mbf1 inhibit frameshifting in a cooperative manner, perhaps due to their interactions with mRNA or to Mbf1’s interaction with the ribosome. First, the role of Rps3 in this process is likely to involve interactions with either the incoming mRNA or proteins external to the ribosome. The *RPS3* mutations that affect frameshifting map to residues (*S104*, *L113, G121*, *K108*) on two α-helices or their connecting loop right next to the entering mRNA. Although this section of Rps3 is involved in helicase activity and initiation selectivity (Dong et al., 2017, Takyar et al., 2005), the residues mutated in frameshifting selections were not specifically those involved in these activities. Instead, these residues all sit on the solvent side of the ribosome and could form an interface interacting with mRNA or mRNA-bound proteins. Moreover, these residues in which mutations affect reading frame maintenance are specifically conserved in eukaryotes (and differ in archaeal and bacterial Rps3), consistent with a eukaryotic-specific mechanism. Second, Mbf1 is likely to interact with either or both of the mRNA and the ribosome, based on work by others (Beckmann et al., 2015, Blombach et al., 2014, Klass et al., 2013, Opitz et al., 2017). Mbf1 is sufficiently abundant with ~85,000 molecules per cell to participate in general translation cycles, although it is less abundant than core ribosomal proteins (~200,000) (Kulak et al., 2014). Moreover, Mbf1 is likely to interact with the ribosome, since the archaeal homolog of Mbf1 weakly associates with the ribosome through its HTH domain and the linker at the N terminus of this domain, which are both conserved with eukaryotes (Blombach et al., 2014). We note that our frameshifting mutations cluster in this region of Mbf1. Intriguingly, the apparent RNA binding domain maps to the less conserved N terminal domain (Klass et al., 2013). It remains to be seen how these activities come together to regulate reading frame.

Frameshifting occurs by a mechanism that involves the interplay between the two adjacent codons, in which I•A wobble interaction in the P site in conjunction with competition between tRNAs entering the A site results in the frameshift, consistent with a model proposed by Baranov *et al*. (Baranov et al., 2004). First, we demonstrated that, in the *asc1*Δ *mbf1*Δ double mutant, ribosomes frameshift at a single CGA-CGG codon pair (in a particular context) when the CGA codon occupies the P site. We infer that CGA codon in the P site is generally important for frameshifting, because six of the seven codon pairs on which ribosomes frameshift are CGA-NNN and the three efficient pairs are CGA-CNN. The wobble interaction between the CGA codon and tRNA could weaken the interaction between mRNA and the ribosome, which in turn could slow down the elongation cycle. Second, we found that frameshifting is influenced by the abundance of the in frame and out of frame tRNAs for next position, which implies that the frameshift occurs after translocation of the CGA from the A site to the P site. We speculate that the flexibility of the wobble base pair interaction between inosine and other nucleotides could actively facilitate the acceptance of out-of-frame A site tRNA. For instance, we consider that a rare instance in which the A base in CGA is bulged out might be stabilized by the very strong I•C interaction, increasing the time available to accept the out-of-frame tRNA.

The eukaryotic specific reading frame maintenance activity, involving Mbf1 and ribosomal proteins Rps3 and Asc1, is likely to be important for translation accuracy in the yeast genome. Mutations in either *RPS3* or *MBF1* suppressed frameshifting mutations in several native yeast genes (Hendrick et al., 2001). Moreover, mutations in *MBF1* and *ASC1* resulted in detectable frameshifting in a set of native gene sequences with only a single inhibitory codon pair flanked by 6 adjacent codons on each side, although it is apparent that the frameshifting potential within a particular sequence is not simply due to the presence of a single inhibitory codon pair. These results confirmed that Mbf1 with Rps3 and Asc1 play a critical role in maintaining the reading frame during normal translation cycles. It is still unknown why this eukaryote-specific reading frame maintenance system evolved and why it is important to eukaryotes, but not bacteria.

## MATERIALS AND METHODS

### Strains, plasmids, and oligonucleotides

Strains, plasmids, and oligonucleotides used in these studies are listed in Supplementary Tables 9-11. Parents for all yeast strains described in this study were either BY4741 (*MAT****a*** *his3*Δ1 *leu2*Δ0 *met15*Δ0 *ura3*Δ0) or BY4742 (*MATα his3*Δ1 *leu2*Δ0 *lys2*Δ0 *ura3*Δ0) (Open Biosystems). The *GLN4*_(1-99)_-(CGA)_6_+1-*URA3* reporter used in the selection was constructed with PCR-amplified DNAs (using oligonucleotides OJYW085, 086, 041, 089, 095 and 099), assembled by Ligation Independent Cloning (LIC) methods (Alexandrov et al., 2004, Aslanidis and de Jong, 1990) and then integrated into the *CAN1*/*YEL063C* locus on the chromosome V, selecting for canavanine-resistance; constructs were checked by sequencing of genomic PCR fragments. RNA-ID reporters were constructed as described previously and integrated at the *ADE2* locus, using selection with *MET15* marker in *MAT****a*** strains or *S.pombe HIS5* marker in *MATα* strains (Dean and Grayhack, 2012, Gamble et al., 2016, Wolf and Grayhack, 2015).

Yeast strains bearing gene deletions (*mbf1*Δ, *slh1*Δ, and *hel2*Δ) were constructed by amplification of the *kan^R^* cassette in the yeast strain from the corresponding knockout strain in the systematic deletion collection (Open Biosystems) (Giaever et al., 2002). The *MAT****a*** yeast strain bearing a deletion of *RPS3* was constructed by amplification of *ble^R^* cassette (Gueldener et al., 2002) (oligos OW443 and OW445) and integration of this DNA into a strain bearing an *URA3* [*RPS3*] covering plasmid (pEAW433). Yeast strains bearing deletions of *ASC1* marked with the *S. pombe HIS5* marker (AW768), which have been described previously (Wolf and Grayhack, 2015), were constructed and maintained in the presence of a plasmid born copy of *ASC1* on a 2μ, *URA3* plasmid. To obtain the *asc1*Δ strain from the selection parent strain, the *ASC1* gene was deleted by a *ble^R^* cassette obtained by PCR amplification with oligos OW125 and OW126.

Plasmids bearing the *MBF1* gene were constructed by amplification of chromosomal *MBF1* gene from −580 in 5’ UTR to +300 in 3’ UTR with oligos OJYW124 and OJYW125, followed by cloning into the 2μ, *LEU2* vector (pAVA0577) and into the *CEN*, *LEU2* vector (pAVA0581) to create pEJYW203 and pEJYW176 respectively. The chromosomal HA-tagged *MBF1* was constructed by PCR amplification of HA-*kan^R^* sequence from pYM45 (Euroscarf) (Janke et al., 2004) with oligos OJYW130 and OJYW132, bearing homology to *MBF1*, followed by integration into the *MBF1* locus. This *MBF1-HA Kan^R^* cassette from −580 in 5’UTR to +300 in 3’UTR of *MBF1* (+1992 including *Kan^R^* sequences) was amplified from the chromosome with oligos OJYW157 and OJYW158, cloned into the XmaI and NheI sites in Bluescript as pEJYW279. The *mbf1* point mutations *K64E* and *I85T* were individually introduced into the plasmid pEJYW279 to make pEJYW302 and pEJYW307 respectively. The *mbf1-K64E* cassette was directly PCR-amplified from the mutant strain YJYW290-P38 with oligos OJYW157 and OJYW158 followed by digestion with XmaI and BamHI and integration into these two sites on pEJYW279. The *mbf1-I85T* mutation was introduced by PCR amplification from *MBF1-HA* cassette with OJYW170, which contains the mutation, and OJYW166, followed by integration into pEJYW279 between BamHI and AatII sites. Reconstructed *mbf1* point mutants were introduced into YJYW2566 (BY4741, *HIS3^+^*) with XmaI/NheI digested pEJYW302 and pEJYW307 selecting with *Kan^R^* marker.

### Selection for frameshifting mutants and identification of mutations

Ura^+^ mutants were selected from 40 independent cultures of each *MAT****a*** and *MATα* parent strains (YJYW289, YJYW329), and then were analyzed by flow cytometry to measure GFP and RFP expression. Ura^+^ GFP^+^ mutants, indicative of increased frameshifting efficiency, were selected for further study, with an emphasis on mutants that exhibited higher levels of frameshifting, i.e., GFP/RFP >4, (28% *MATα* and 66% *MAT****a*** mutants). Diploids between 12 *MAT****a*** mutant and 20 *MATα* mutants were created by mating in YPD for 2 hours at 25 °C and selection on SD-Lys-Leu-His media for diploid cells, followed by streaking for single colonies. Then overnights of the resultant diploids and their haploid parents were spotted on SD-Leu and SD-Leu-Ura plates, which were grown at 30 °C.

To identify the relevant mutation in YJYW290-P25, we obtained the Leu^-^ derivative of this mutant (YJYW315) by screening replica plated single colonies from an overnight in YPD on YPD and SD-Leu plates. The Ura^+^/FOA-sensitive phenotype of this mutant was complemented with a genomic tiled library (Jones et al., 2008), selecting for FOA-resistant cells. First, 17 pools of DNA, each of which contained 96 plasmids (Jones et al., 2008), were transformed individually with >1000 colonies per plate. Transformants of each pool were then scraped and saved in 2 ml YPD+8 % DMSO. These saves were plated based on their OD_600_ (2×10^7^ cells/OD_600_ x ml) to obtain approximately 5,000 cells on SD-Leu and 50,000 cells on SD-Leu+0.5xFOA. For 16 of 17 pools, there were no colonies on the FOA plates, while transformants of pool 15 had 330 FOA-resistant colonies with 1404 colonies on SD–Leu plate, corresponding to FOA-resistance for 2.3% cells. The plasmids responsible for FOA-resistance was identified by complementing with plasmids from individual rows and columns in this pool as described above, followed by complementation with individual plasmids. Two plasmids from this pool conferred FOA-resistance and share a single gene, *MBF1*. The *MBF1* gene in 19 recessive mutants was amplified from their genomic DNA with oligos OJYW124 and OJYW125, followed by sequencing to confirm the mutated residues.

Whole genome sequencing on two dominant *MATα* mutants was performed to identify the mutated genes. For each strain, ~30 OD_600_ yeast cells were harvested and re-suspended in 1 ml prep buffer (2% Triton X-100, 1% SDS, 100 mM NaCl, 10 mM Tris-Cl pH 8.0, 1 mM EDTA) with ~1.5 g Zirconia/Silica beads (from BioSpec, catalog# 11079105z) and 1 ml PCA pH 8.0. The suspension was then vortexed at top speed for 3 minutes and mixed with 1 ml TE pH 8.0, followed by centrifugation in prespun PLG tubes (from 5prime, catalog# 2302830). Nucleic acids in the aqueous layer were ethanol precipitated with 5 ml 100% ethanol, followed by freezing on dry ice and centrifugation for 20 minutes at 4,000 rpm at 4 °C. The pellet was re-suspended in 200 μl TE and incubated at room temperature for 1 hour with 0.2 μg/μl RNaseA to remove RNA contamination, followed by addition of 200 μl 1 M Tris-Cl pH 8.0, 2 μl of 5 mg/ml glycogen and 400 μl PCA, and centrifugation for 2 minutes at top speed at 4 °C. The aqueous layer (~360 μl) was precipitated with 720 μl 100% ethanol and frozen on dry ice for 15 minutes; resulting pellets were re-suspended in 100 μl TE pH 8.0 and 100 μl 1 M Tris-Cl pH 8.0, followed by precipitation again with 400 μl 100% ethanol. The DNA pellet was then washed with 500μl 70% ethanol and finally re-suspended in 50 μl sterile ddH_2_O. Whole genome sequencing was performed by the UR Genomics Research Center resulting in *RPS3* mutations in these two *MATα* mutants. Mutations in two *MAT****a*** dominant mutants were then identified by amplification of *RPS3* cassette with oligos OJYW159 and OJYW210, followed by sequencing.

### Analysis of yeast growth

Appropriate control strains (previously studied) and 2-4 independent isolates of each strain being tested were grown overnight at 30°C in media indicated, diluted to obtain OD_600_ of 0.5, then serially diluted 10-fold twice; 2 μl diluted cells were then spotted onto the indicated plates and grown at different temperatures for at least two days.

### Flow cytometry

To examine mutants in either *RPS3* or *ASC1*, reporters were introduced into sets of strains bearing an *URA3* covering plasmid with either *RPS3* or *ASC1*, depending upon the chromosomal deletion. All sets of strains in a given panel contained the same *URA3* plasmid. Prior to analysis of GFP expression, strains were streaked on FOA containing plates, then single colonies were grown for analysis by flow cytometry.

Yeast strains bearing the modified RNA-ID reporters were grown overnight at 30 °C in YP media (for strains without plasmid) or appropriate synthetic drop-out media (for strains with plasmid) containing 2% raffinose + 2% galactose + 80 mg/L Ade. The cell culture was diluted in the morning such that to the culture had a final OD_600_ between 0.8-1.0. Analytical flow cytometry and downstream analysis were performed for 4 independent isolates of each strain (Outliers were rejected using a Q test with >90% confidence level) as previously described (Dean and Grayhack, 2012). Each flow experiment was also performed with proper controls including a GFP^-^, RFP^+^ strain. The GFP/RFP value from this control strain was subtracted from all tested strains on the same day to show signals above background (negative values are set to 0). P values were calculated using a one-tailed or two-tailed homoscedastic t test in Excel, as indicated in Supplementary Table 12.

### Western blotting

Western analysis of the GFP fusion proteins in the modified RNA-ID reporter and Mbf1 protein in yeast strains were performed with anti-HA antibody as described previously (Gelperin et al., 2005).

### RT-qPCR

The GFP mRNA measurement with reverse transcription (RT) reaction and quantitative PCR was performed as described previously (Gamble et al., 2016). For each tested strain, three biological replicates were analyzed, while one of the isolates in each experiment was performed with two technical replicates to obtain standard curve. P values were calculated using a one-tailed homoscedastic t test in Excel, as indicated in Supplementary Table 12.

### Purification of frameshifted peptide

To purify the frameshifted peptide from yeast, a *LEU2* plasmid containing either in658 frame or +1 frame protein purification constructs were transformed into the *asc1*Δ *mbf1*Δ strain (YJYW378). Two independent transformants (FOA treated) of each construct were grown overnight in SD-Leu media and transferred into 80 ml S-Leu+2% raffinose media in the morning. After reaching an OD_600_ of 0.8-1.2, expression of the GST-StrepII-ZZ construct was induced by addition of 40 ml 3xYP+6% galactose and growth was continued for 10 hours. Cells were collected by centrifugation and cell pellets were quick frozen on dry ice. The cell pellets were re-suspended in 1 ml extraction buffer (50 mM Tris-Cl pH 7.5, 1 mM EDTA, 4 mM MgCl_2_, 5 mM DTT, 10% Glycerol, 1 M NaCl, 2.5 μg/ml leupeptin, 2.5 μg/ml pepstatin) and lysed with bead beating (10 repeats of 20 second beating followed by 1 minute on ice), essentially as described previously (Quartley et al., 2009). The cell lysate was collected from the bead beating tubes by puncturing the bottom with a hot needle and blowing with low pressure air. Solid contents were removed by centrifugation before the remaining lysate was divided into half and purified on either GSH or Streptactin resin.

For GST purification: the cell lysate was first diluted with equal volume No Salt Wash Buffer (50 mM Tris-Cl pH 7.5, 4 mM MgCl_2_, 5 mM DTT, 10% Glycerol) to bring the salt to 0.5 M NaCl. GSH resin [Glutathione sepharose-4B from GE, catalog# 17-0756-01; pre-washed with Wash Buffer (No Salt Wash Buffer + 0.5 M NaCl)] (50 μl/ ml of lysate) was added to the diluted cell lysate and the mixture was nutated for 3 hours at 4 °C. The resin was separated from the liquid by centrifugation at low speed (<3,000 rpm) and washed twice with 0.5 ml Wash Buffer followed by 20-minute nutation. The bound protein products were then eluted by nutating for 40 min with 100 μl Elution Buffer (Wash buffer + 20 mM NaOH + 25 mM glutathione); the elution step was repeated to increase the yield.

For Strep purification: the cell lysate was diluted with 5x volumes No Salt Wash Buffer (100 mM Tris-Cl pH 7.5, 1 mM EDTA, 2.5 μg/ml leupeptin, 2.5 μg/ml pepstatin) to bring the salt to 150 mM NaCl. MagStrep “type3” XT beads [from IBA, cat# 2-4090-002; pre-washed with Wash Buffer (No Salt Wash Buffer + 150mM NaCl)] were added to the diluted cell lysate (80 μl/ 3.3 ml diluted cell lysate). After nutating for 2 hours at 4 °C, resins were separated from liquid using a magnetic separator, then the resin was washed with 1 ml Wash Buffer three times without additional incubation. The bound protein products were then eluted by adding 50 μl Elution Buffer (Wash buffer + 50 mM biotin) and nutating for 10 minutes, followed by separation using magnetic separator; the elution step was repeated to increase the yield.

### Mass spectrometry

The elution samples from both GST and Strep purification were analyzed by SDS-PAGE, followed by staining with Coomassie Blue. The bands from Strep purification of both in-frame and +1 frame constructs were excised and analyzed on the Q Exactive Plus Mass Spectrometer in the Mass Spectrometry Resource Center of the University of Rochester Medical Center.

## SUPPLEMENTAL INFORMATION

Supplemental Information includes 12 supplementary tables provided in 2 supplementary files.

## AUTHOR CONTRIBUTIONS

J.W. and E.G. wrote the manuscript. J.W. performed most experiments while Q.Y. examined effect of overproduction of Mbf1 on frameshifting in *RPS3* mutants and J.Z. examined frameshifting from native sequences.

## ACKNOWLEDGEMENTS

We thank Eric Phizicky, Christina Brule, Andrew Wolf and Lu Han for discussions of the science and comments on the manuscript; Christina Brule and Blake Bentley for assistance with experiments. This research has been facilitated by the services and resources provided by the University of Rochester Mass Spectrometry Resource Laboratory and NIH instrument grant (1S10OD021486-01). We thank Genomics Research Center for performing high-throughput sequencing library construction, sequencing, and primary data analysis for this study. We also thank the URMC Flow Cytometry Resource for technical support. This work was supported by NIH grant R01 GM118386 to E.J.G.

---

**Figure 1- figure supplement 1.**
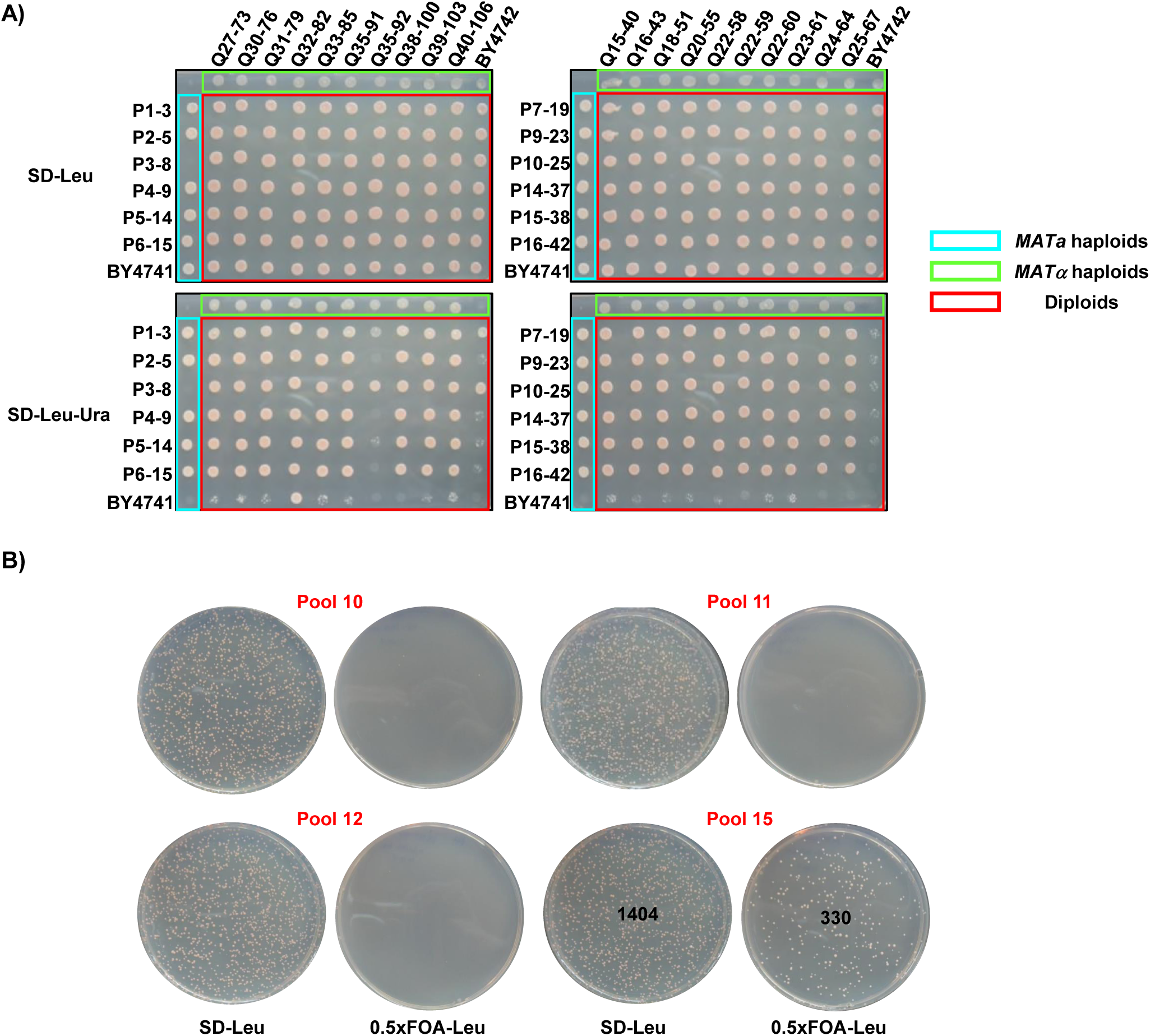
Classification of dominant and recessive mutations and complementation of a recessive mutation. **(A)** Analysis of complementation and dominant/recessive nature of mutations. Twelve *MAT****a*** mutants were crossed with 20 *MATα* mutants, as well as with their selection parents. An Ura^+^ phenotype of resulting diploids with the wild type parent indicated that 3 mutants were dominant while the Ura^+^ phenotype of mutants crossed with each other indicated one major complementation group among recessive mutants. **(B)** Introduction of the Prelich library pool 15 DNA resulted in FOA-resistant cells (Ura^-^) which indicates suppression of the frameshifting phenotype.

**Figure 1- figure supplement 2.**
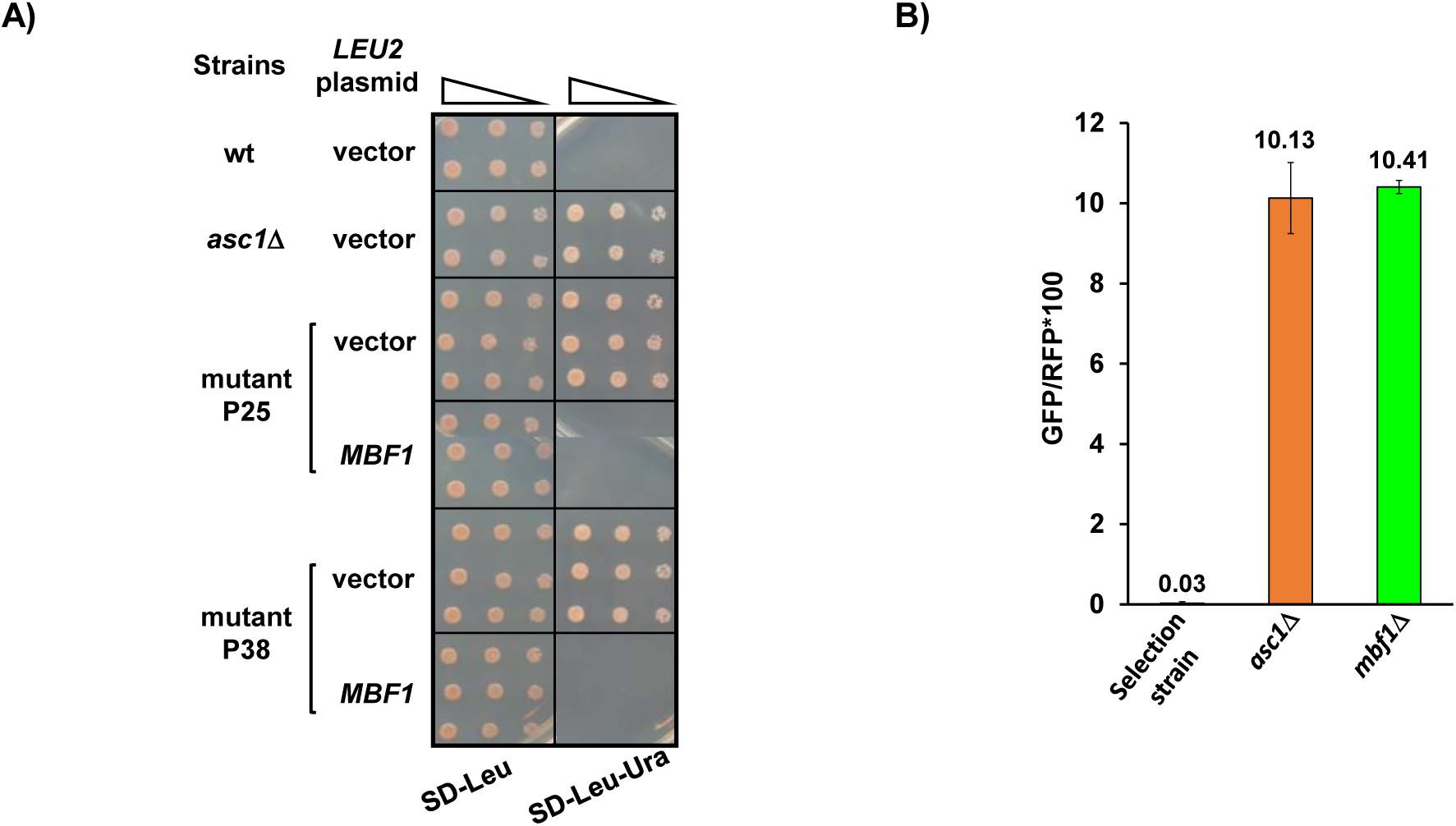
Confirmation that mutations in *MBF1* are responsible for frameshifting. **(A)** Plasmid-borne *MBF1* gene suppressed the Ura^+^ phenotype of mutants P25 and P38. **(B)** Deletion of the *MBF1* coding sequence in the parent GFP^-^ strain resulted in GFP^+^ phenotype.

**Figure 1- figure supplement 3.**
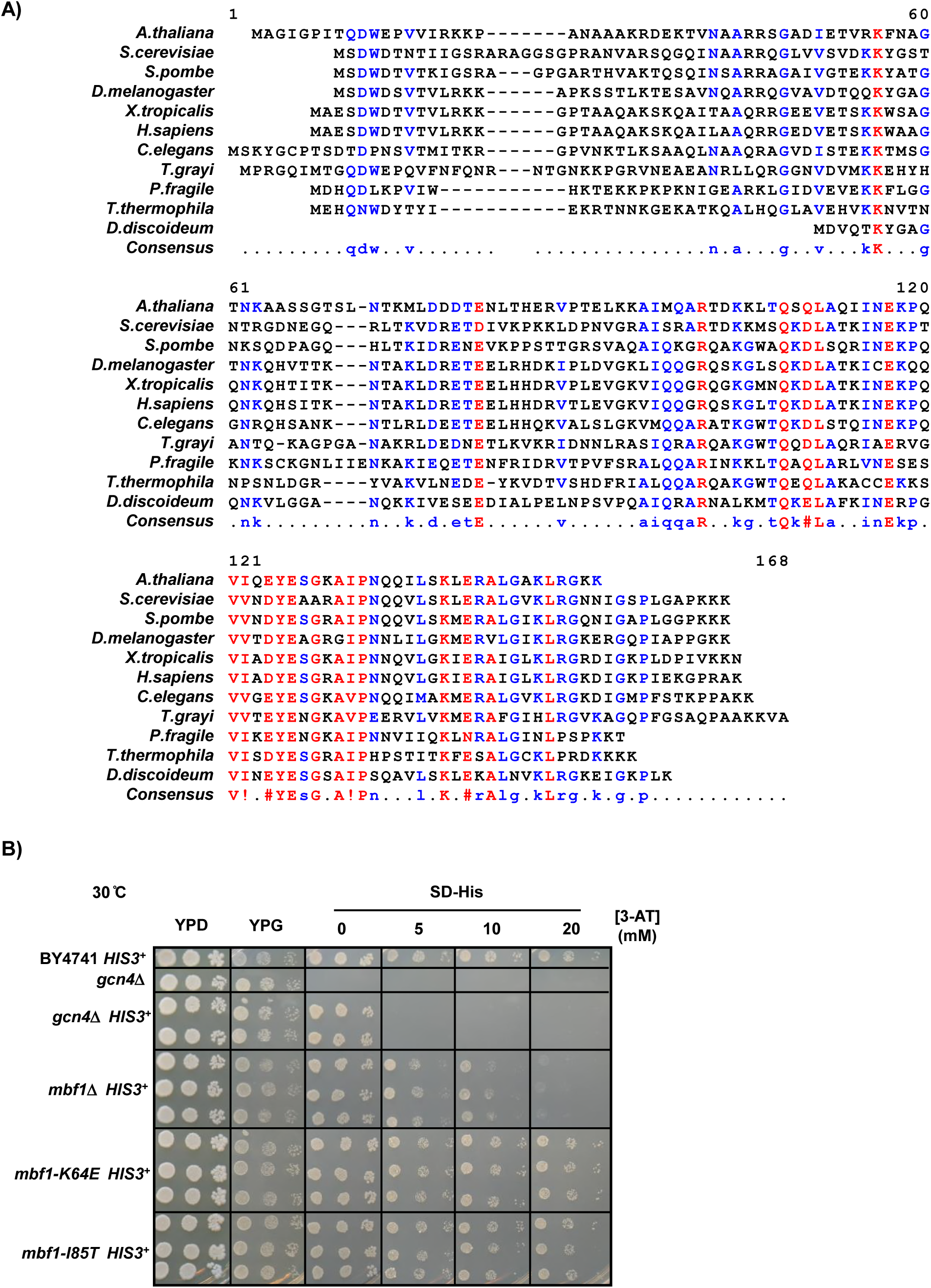
Mbf1 is conserved and frameshifting mutations do not exhibit sensitivity to 3-AT. **(A)** Amino acid sequence alignment of Mbf1 protein from 11 eukaryotic species using MultAlin (http://multalin.toulouse.inra.fr/multalin/) (Corpet, 1988). The color of markers corresponds to the consensus level of this residue (Blue: 50%-90%, Red: 90%) **(B)** Frameshifting *mbf1*-*K64E* and *I85T* mutants grow like wild type on plates with 3-aminotriazole and do not display a *gcn4*Δ phenotype. The *mbf1*Δ strains are more resistant to 3-AT than *gcn4*Δ strains.

**Figure 3- figure supplement 1.**
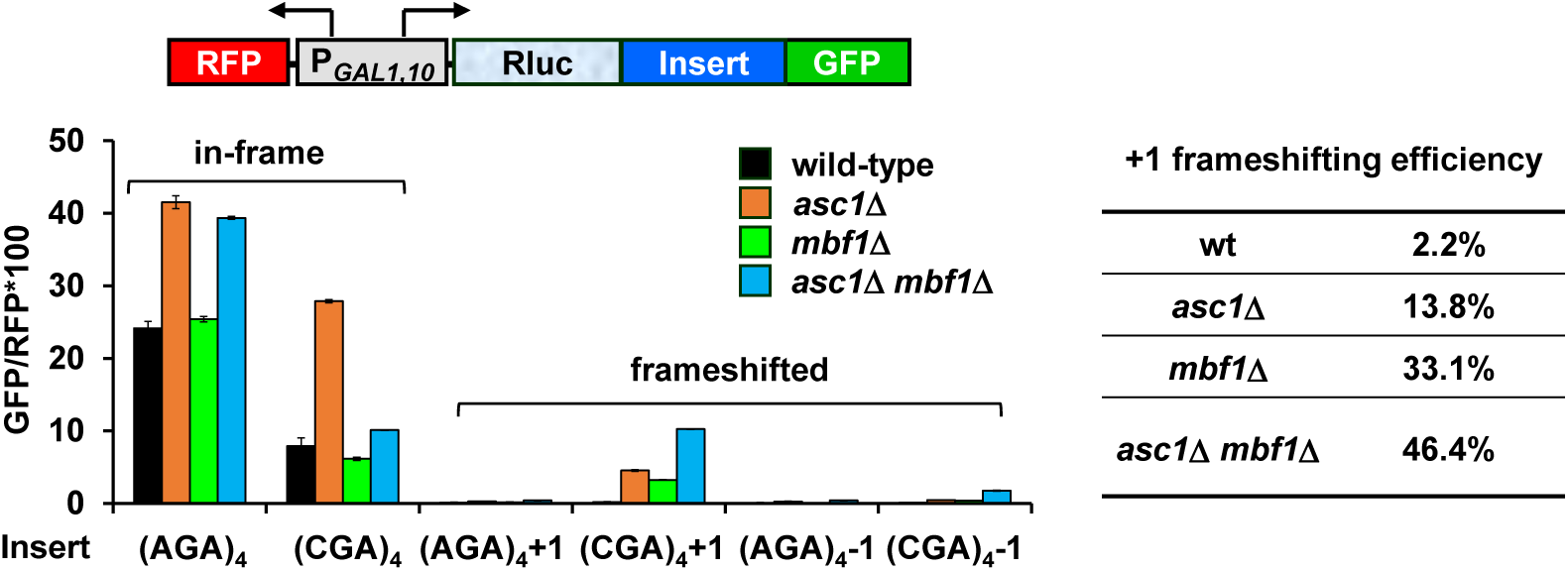
Analysis of effects of *asc1*Δ, *mbf1*Δ *and asc1*Δ *mbf1*Δ mutations on expression of Rluc-GFP reporters containing four adjacent Arg codons (AGA versus CGA) in 0, +1, and −1 reading frames. Mutation of either *ASC1* or *MBF1* allows frameshifting in the (CGA)_4_+1 reporter, and mutation of both *ASC1* and *MBF1* results in significantly more frameshifted GFP/RFP. The +1 frameshifting efficiency [(+1 GFP/RFP) / (+1 GFP/RFP + in-frame GFP/RFP + −1 GFP/RFP)] of all four strains is shown in the table.

**Figure 3- figure supplement 2.**
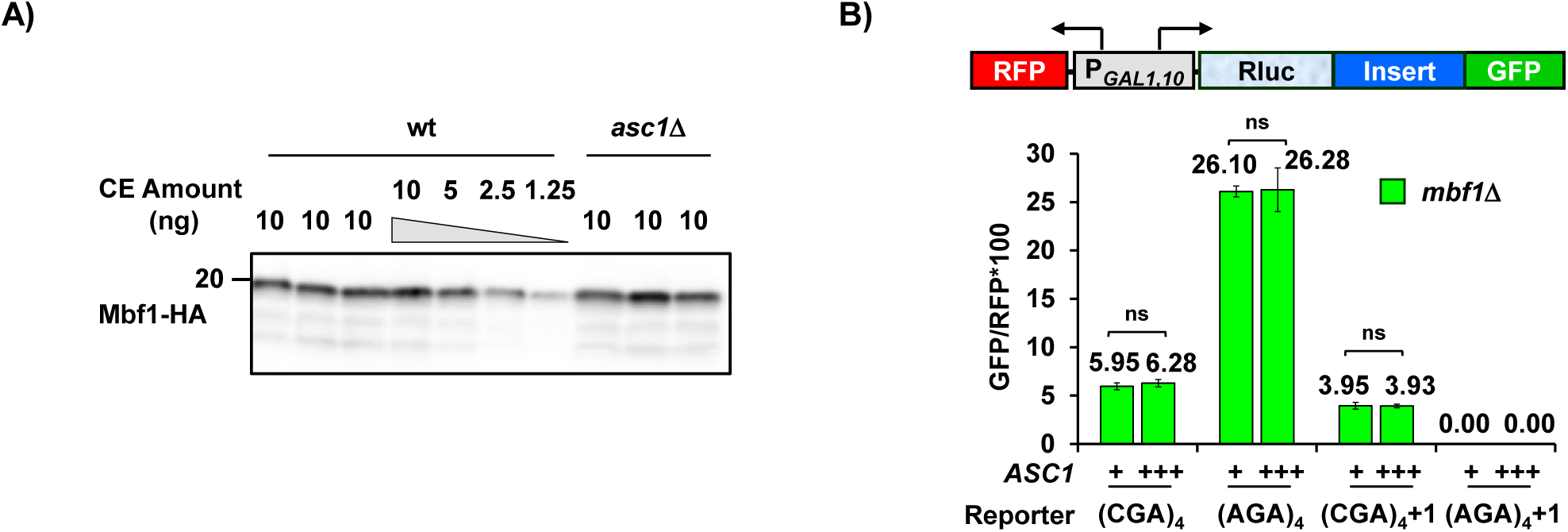
Frameshifting is likely not due to reduction of Mbf1 protein in *asc1*Δ mutant nor to limiting Asc1 protein in *mbf1*Δ mutant. (**A)** Western analysis of HA tagged Mbf1 in the *asc1*Δ mutant (3 independent isolates shown) compared to the wild-type strain (4 independent isolates shown) indicates that Mbf1 levels were similar in both strains. **(B)** Overexpression of Asc1 does not affect either in-frame read-through or frameshifting at CGA codon repeats in the *asc1*Δ strain. ns: p>0.05.

**Figure 4- figure supplement 1.**
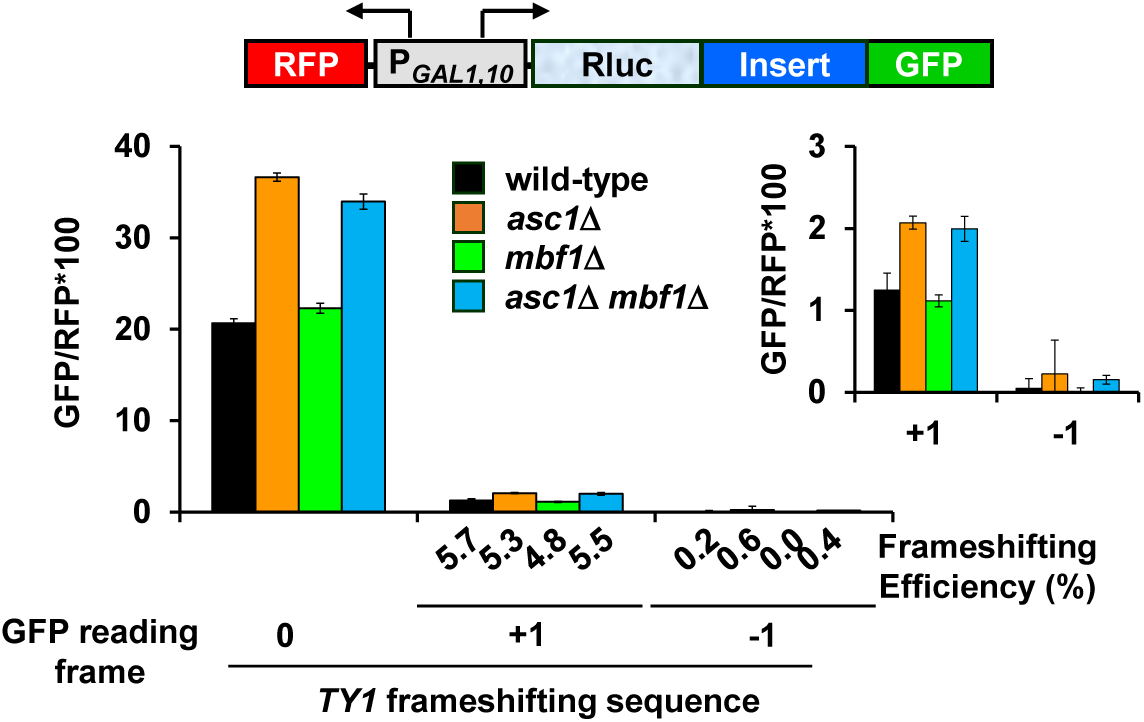
Deletion of *MBF1* and/or *ASC1* does not affect efficiency of programmed frameshifting in the *TY1* transposon. Analysis of effects of *asc1*Δ*, mbf1*Δ *and asc1*Δ *mbf1*Δ mutations on expression of *GLN4*-GFP reporters containing the yeast *TY1* programmed frameshift site (Belcourt and Farabaugh, 1990).

**Figure 5- figure supplement 1.**
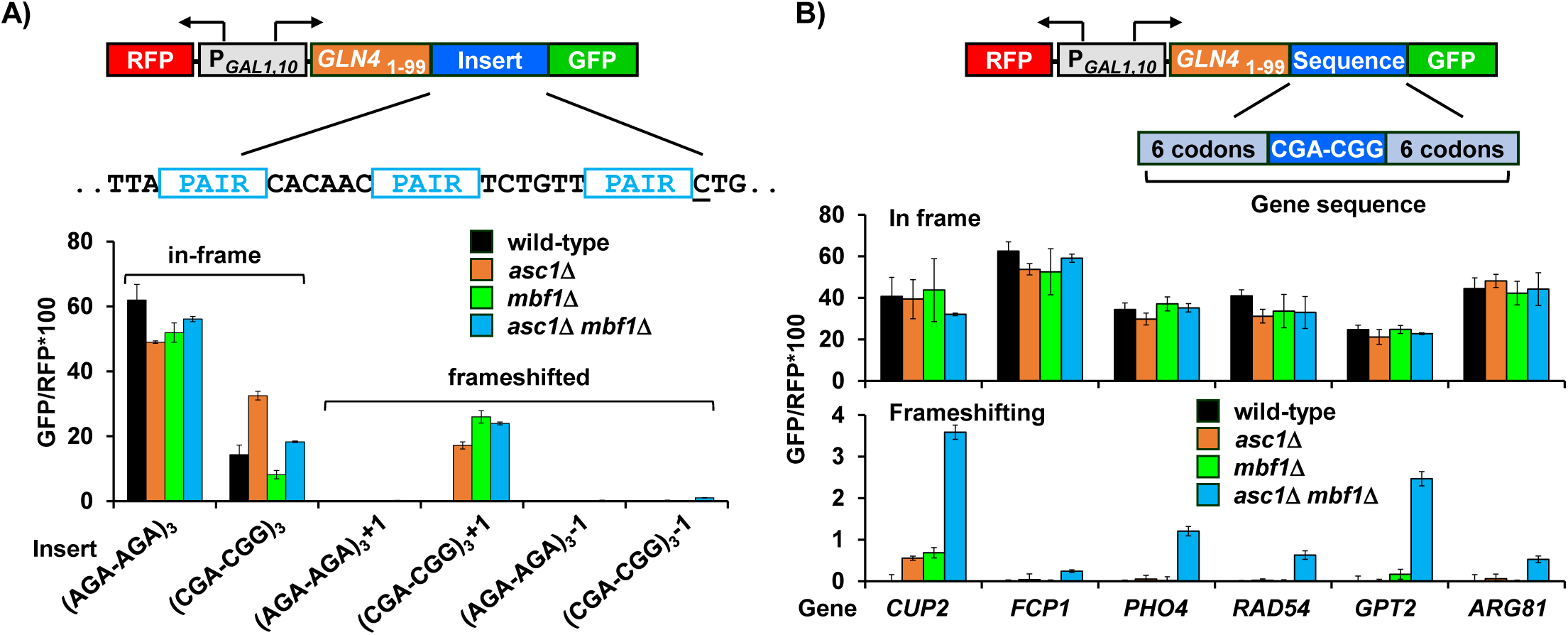
Analysis of frameshifting at CGA-CGG codon pairs. **(A)** Analysis of effects of *asc1*Δ*, mbf1*Δ *and asc1*Δ *mbf1*Δ mutations on expression of *GLN4*_(1-99)_-GFP reporters containing three Arg-Arg codon pairs (AGA-AGA versus CGA-CGG) in 0, +1, and −1 reading frames. In this RNA-ID reporter, CAC is the 3’ codon downstream of the first codon pair. Mutation of either *ASC1* or *MBF1* alone allows extremely efficient frameshifting in this (CGA-CGG)_3_+1 reporter, and mutation of both *ASC1* and *MBF1* does not result in significantly more frameshifted GFP/RFP. **(B)** Analysis of effects of native yeast gene sequences containing a single CGA-CGG codon pair on in-frame and frameshifted expression of GFP. In each case, six codons upstream and downstream of the CGA-CGG were inserted into the *GLN4*_(1-99)_-GFP reporter in frame and with a +1 frameshift after the inserted sequence. Expression of GFP/RFP was measured in wild type, *asc1*Δ*, mbf1*Δ *and asc1*Δ *mbf1*Δ mutants. These native yeast sequences can provoke detectable frameshifting in the *asc1*Δ *mbf1*Δ strain without largely affecting in-frame read-through.

